# The constitutive androstane receptor protects against metabolic dysfunction-associated steatotic liver disease through sex specific interactions with the gut microbiota

**DOI:** 10.64898/2026.10.08.757558

**Authors:** Marine Huillet, Céline M.P. Martin, Guillaume Magne, Frédéric Lasserre, Justine Bruse, Valérie Alquier-Bacquié, Arnaud Polizzi, Clémence Rives, Prunelle Perrier, Noémie Poirier-Jaouen, Sarra Smati, Géraldine Garcia, Jelskey Gomes, Elodie Rousseau-Bacquié, Claire Naylies, Gaëlle Payros, Yannick Lippi, Théo Perion, Emilien L. Jamin, Rolle-Kampczyk Ulrike, Martin von Bergen, Justine Bertrand-Michel, Véronique Pons, Alexandra Montagner, Pierre Gourdy, JingHong Wan, Sophie Lotersztajn, Emmanuelle Vallez, Anne Muhr-Tailleux, Bart Staels, Charlène J.G. Dauriat, Benoit Chassaing, Michael Schupp, Yuna Blum, Anne Fougerat, Laurence Gamet-Payrastre, Bruno Lamas, Hervé Guillou, Nicolas Loiseau, Sandrine Ellero-Simatos

## Abstract

**Background:** The gut-liver axis is involved in Metabolic dysfunction-Associated Steatotic Liver Disease (MASLD), but the host molecular actors involved in such dialogue have not yet been fully characterized.

**Objective:** We investigated the sex-dependent role of the Constitutive Androstane Receptor (CAR) as a hepatic sensor of microbial signals in MASLD pathophysiology.

**Design:** Male and female *Car+/+* and *Car-/-* mice were fed a western diet (WD) for 4, 14 or 35 weeks. We performed plasma biochemistry, liver histology, bile acid and cholesterol quantification, untargeted multi-omics approach combining liver RNA-sequencing and 16S rRNA sequencing, together with co-housing, fecal microbiota transplantation and analysis of publicly available transcriptomics data in human cohorts.

**Results:** Short-term WD revealed CAR-dependent protection against steatosis, together with perturbations in the transcriptional signatures of multiple transcription factors involved in hepatocyte identity in males. After 14 weeks, CAR deletion increased WD-induced liver damage, inflammation and fibrosis, and disrupted bile acid, cholesterol and gut microbiota homeostasis, in males only. Co-housing and fecal microbiota transplantations revealed that a *Car+/+* gut microbiota protected against WD-induced liver damage, while a *Car-/-* microbiota was reshaped by a *Car+/+* host. No CAR-dependent phenotype was observed in females. Finally, in human cohorts, hepatic CAR expression decreased in advanced clinical stages of MASLD.

**Conclusion:** CAR is a sex-dependent protective nuclear receptor against MASLD progression, acting both as a transcriptional regulator of hepatocyte identity and as a mediator of the host-microbiota crosstalk.

**KEY MESSAGES:** *What is already known on this topic:*

- The gut-liver axis contributes to Metabolic dysfunction-Associated Steatotic Liver Disease (MASLD) progression.
- CAR is a liver enriched nuclear receptor with sex-dependent activity, recognizing a broad range of xenobiotic and endobiotic metabolites and regulating xenobiotic, glucose, lipid, bile acid and cholesterol metabolism. New studies have also unveiled its ability to recognize bacterial-derived metabolites under physiological conditions. However, its role in MASLD remains controversial.
- The only FDA-approved drug for advanced MASLD targets a liver nuclear receptor (thyroid hormone receptor-β) but is effective in fewer than a third of patients.

*What this study adds:*

- CAR is required for the adaptive transcriptional response of hepatocytes to a western diet, acting as an upstream regulator of multiple transcription factors during early MASLD.
- CAR protects against MASLD in a sex-dependent manner through a causal, bidirectional crosstalk with the gut microbiota, as demonstrated by co-housing and fecal microbiota transplantation experiments
- In patient cohorts, hepatic CAR mRNA expression decreases at advanced MASLD clinical stages.

*How this study might affect research, practice or policy:*

- Nuclear receptor mechanisms in metabolic disease should be systematically assessed for sex dimorphism, as protective pathways identified in one sex may not generalise to the other.
- These findings suggest that both the gut microbiota and CAR itself could be targeted, in a sex-informed manner, as therapeutic strategies for MASLD.

**VISUAL ABSTRACT:** 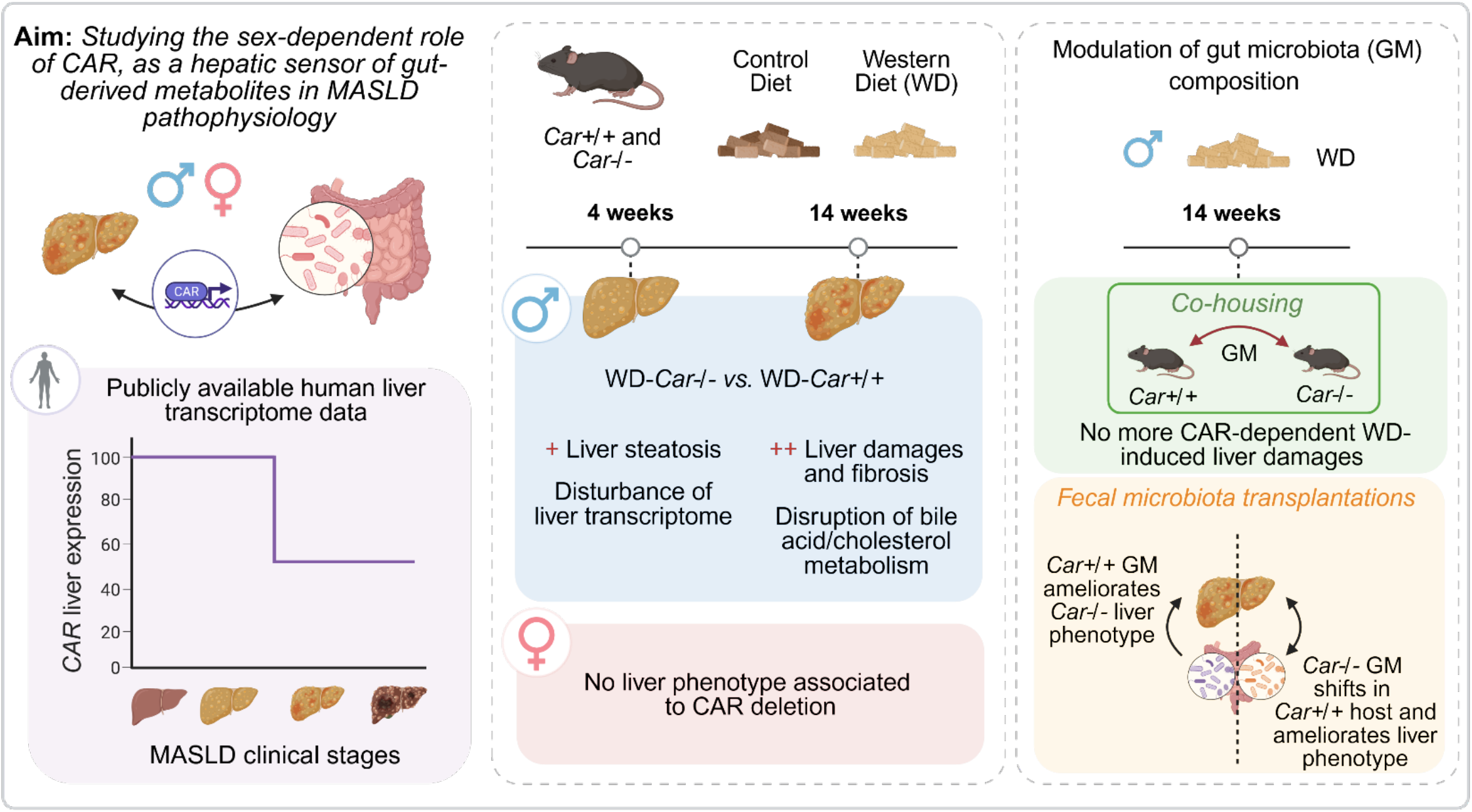

## INTRODUCTION

Metabolic dysfunction-Associated Steatotic Liver Disease (MASLD) is the most common chronic liver disease worldwide. MASLD is a multifactorial disease associated with metabolic disorders such as obesity and type 2 diabetes and is thought to affect at least one quarter of the world population^1,2^. MASLD is characterized by a broad spectrum of liver damages ranging from benign steatosis to Metabolic dysfunction-Associated SteatoHepatitis (MASH), which can further progress to fibrosis, cirrhosis or hepatocellular carcinoma^3^.

Epidemiological studies have revealed that premenopausal women are protected against MASLD independently of cardiovascular risk factors and the presence of metabolic syndrome^4,5^. This protection is linked to the possible role of estrogens since it fades after the menopause^6–8^. In addition, pre-clinical studies have shown that ovariectomy promotes MASH pathogenesis upon a High Fat Diet (HFD)^9^. However, clinical studies of hormone replacement therapy have not yet formally established exogenous estrogens as liver-protective drugs, despite evidence of their positive effects in pre-clinical studies^10^. Thus, the potential sexually dimorphic mechanisms leading to female protection from MASLD remain unclear^11–13^.

Moreover, the gut microbiota has recently evolved as a new player in the pathophysiology of MASLD^14,15^. Indeed, human studies have identified microbiota alterations in MASLD and found consistent microbiome signatures discriminating healthy individuals from those with steatosis, MASH or cirrhosis^15^. In pre-clinical studies, germ-free and antibiotic-treated mice are protected from HFD-induced hepatic steatosis^16^. Finally, Fecal Microbiota Transplantation (FMT) experiments from MASLD/MASH patients into germ-free or conventional mice potentiated HFD-induced hepatic steatosis, revealing the causal role of MASLD-dependent gut microbiota dysbiosis on hepatic phenotype^17–19^. The hepatic receptors involved in the gut-microbiota-liver interactions have yet to be characterized. Members of the nuclear receptor superfamily are natural candidates, as they act as ligand-dependent transcription factors, promptly adapting hepatic gene expression and thereafter metabolism upon sensing of fluctuating levels of nutrients, xenobiotics and/or microbial metabolites^20^. Nuclear receptors serve as targets for 16% of approved small molecule drugs^21^ and have the potential to serve as pharmacological targets for MASLD/MASH^22^. In March 2024, resmetirom, a liver-directed thyroid hormone receptor-β agonist has received approval from the FDA for the treatment of MASH and remains the only approved drug for these pathologies^23^. Nevertheless, resmetirom was found effective in less than a third of individuals, potentially due to strong disease heterogeneity^24^. Thus, better understanding of other nuclear receptors’ biology has the potential to enhance potential MASLD/MASH treatments.

We have previously described the Pregnane X Receptor (PXR, official gene symbol NR1I2), a xenobiotic-sensing nuclear receptor, as a hepatic effector of microbiota-derived signals that regulate the host’s lipid metabolism in a sexually dimorphic manner^25^. In the present study, we investigated the potential role of another xenobiotic nuclear receptor, the Constitutive Androstane Receptor (CAR, official gene symbol NR1I3). PXR and CAR were originally characterized as regulators of xenobiotic clearance^26,27^, but many studies have now highlighted their role in numerous metabolic pathways relevant for MASLD progression such as *de novo* lipogenesis, β-oxidation, bile acid metabolism and oxidative stress^28–32^. Interestingly, CAR’s transcriptional activity in the liver is dependent on gut microbiota signals^33^ and is sexually dimorphic^34,35^. Here, we aimed to test the hypothesis that CAR integrates dietary and microbial signals to modulate MASLD progression. We used *Car^+/+^* and *Car^-/-^* mice (thereafter named *Car+/+* and *Car-/-* respectively) exposed to different nutritional interventions and found that CAR protects from MASLD development in a sexually-dimorphic way, through complex bidirectional interactions between host genetics and its gut microbiota. Finally, we observed a significant decrease of CAR mRNA expression during MASLD progression in human clinical cohorts.

## METHODS

See additional methods.

## RESULTS

### CAR buffers the early transcriptional adaptation to WD in the male liver

To investigate the potential role of CAR in steatosis, we investigated the effects of a short-term western-diet (WD) in *Car+/+ vs. Car-/-* mice (figure 1A). After 4 weeks of WD, *Car-/-* males had higher liver weight and steatosis, while lipid accumulation was still minor in *Car+/+* males (figure 1B-E). In females, no significant genotype*diet interactions were observed (figure 1B&C, F&G). Given that CAR functions directly at the genome to control transcription, we performed hepatic RNA-sequencing. This revealed more than 2 times more WD-responsive differentially expressed genes (DEGs) in *Car-/-* males (480 DEGs) than in *Car+/+* males (200 DEGs) (figure 1H&I), and 2 times more DEGs in *Car-/-* females (196 DEGs) than in *Car+/+* females (97 DEGs) (supplemental figure 1), demonstrating that CAR buffers the early transcriptional response to a WD in mouse liver. The most consistently perturbed pathway upon WD was cholesterol metabolism, in both males (figure 1H&I) and females (supplemental figure 1A&B), which is consistent with high liver cholesterol content in WD. However, WD-induced perturbations in the expression of genes involved in cholesterol metabolism were similar between *Car+/+* and *Car-/-* males (supplemental figure 2). Genes specifically regulated in *Car-/-,* and not *Car+/+,* males, were involved in TNFα production and fatty acid metabolism, consistent with increased steatosis and early inflammation in those mice (figure 1H). We next performed an exhaustive transcription factor enrichment analysis (figure 1J). In all groups, we observed significant enrichment in PPARα, PPARγ and NR1H3 (encoding LXR) target genes in WD-compared to chow diet (CD)-fed mice, with positive scores representing an enrichement in the target genes of these transcription factors among genes upregulated upon WD, in agreement with activation of lipid catabolism and anabolism in the early stages of steatosis. Many transcription factor target genes were also enriched among genes down-regulated upon WD in all groups, among which target genes of HNF1α, NR1H4 (encoding FXR), FOXA2 and SREBP1. Strikingly, for these transcription factors, enrichment scores were slightly, but consistenly more negative in *Car-/-* than in *Car+/+* males. In accordance, the largest differences between *Car+/+* and *Car-/-* males all followed the same pattern with 8 transcription factors with positive scores in *Car+/+* males that changed directionality in WD-*Car-/-* males, showing that these transcription factors were activated upon WD in *Car+/+* males and suppressed in *Car-/-* mice. Among these are transcription factors with pleiotropic effects on the hepatocyte identity such as STAT5a, STAT3 and HNF4α.

**Figure 1:**
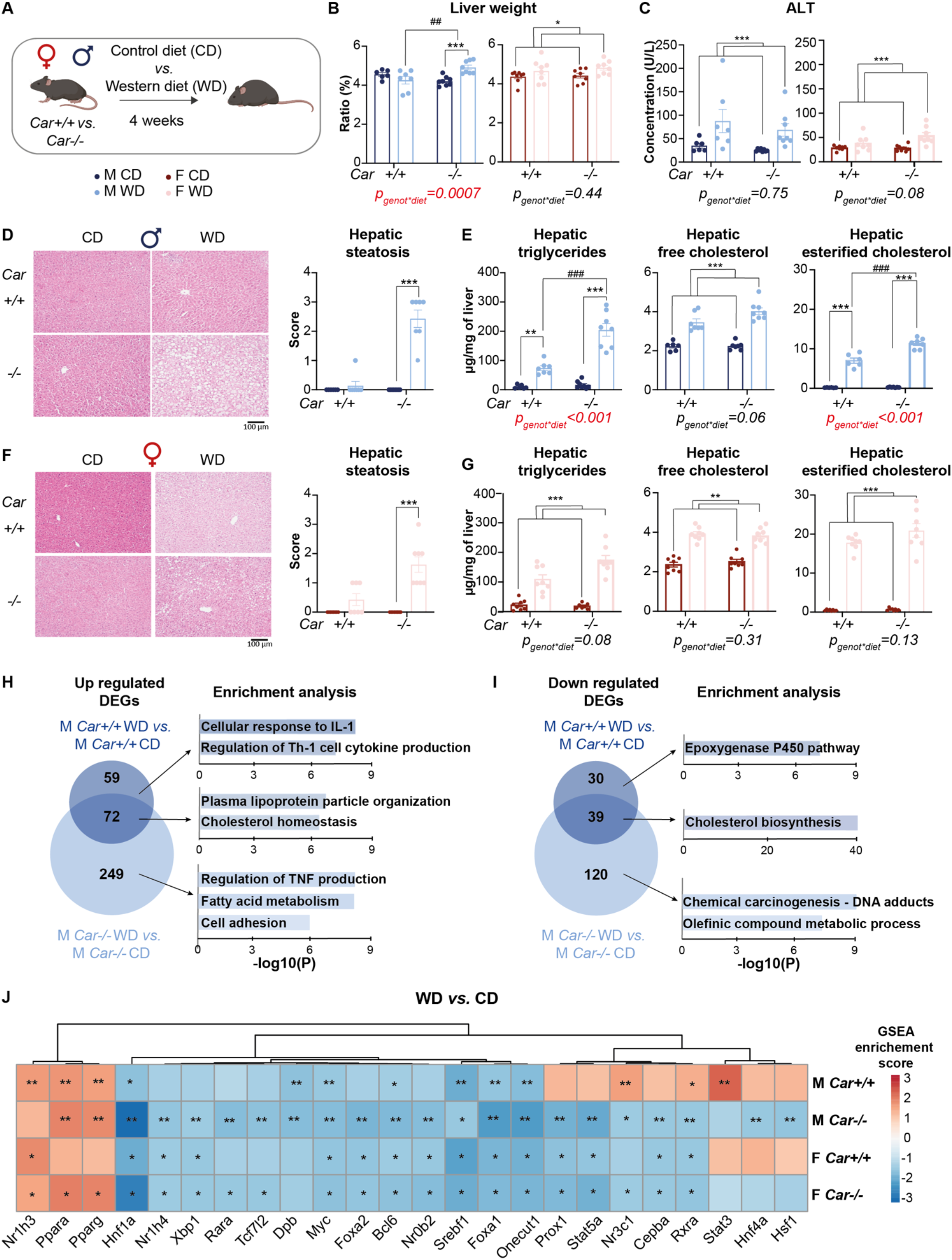
CAR protects male mice from WD-induced steatosis. A) Experimental design. B) Relative liver weights. C) Plasma alanine aminotransferase (ALT) concentrations. D) Representative histological sections of male liver stained with haematoxylin and eosin. Magnification x100. Liver steatosis scores estimated on histological male sections. Scoring: parenchymal involvement by steatosis <5%, 0; 6-33%, 1; 34-66%, 2; >67%, 3. E) Male hepatic triglycerides, free cholesterol and cholesterol esters analysed by gas-liquid chromatography. F) Representative histological sections of female liver stained with haematoxylin and eosin. Magnification x100. Liver steatosis scores estimated on histological male sections. Scoring: parenchymal involvement by steatosis <5%, 0; 6-33%, 1; 34-66%, 2; >67%, 3. G) Female hepatic triglycerides, free cholesterol and cholesterol esters analysed by gas-liquid chromatography. H) Venn diagram representing the number of genes significantly upregulated by WD in Car+/+ and Car-/- male mice (FC>2, p_adj_<0.05) and pathway enrichment analysis. I) Venn diagram representing the number of genes significantly downregulated by WD in Car+/+ and Car-/- male mice (FC<-2, p_adj_<0.05) and pathway enrichment analysis. J) GSEA analysis of transcription factor target genes enrichment analysis in WD- vs CD-fed mice. Only transcription factor with at least one FDR<0.05 were selected for heatmap representation. * FDR<0.1, *\*\* FDR<0.05, *** FDR<0.01. Results are given as the mean ± SEM. *diet effect, #genotype effect. * or # p<0.05, ** or ## p<0.01, *** or ### p<0.001 (two-way ANOVA). Histological scores were analysed by a non-parametric test (Kruskal-Wallis)*.

We wondered whether this massive transcriptional regulation was specific to any liver cell type. Cell type enrichement analysis from DEGs in WD- vs CD-fed males revealed that down-regulated genes originated mostly from hepatocyte progenitors and mature hepatocytes, both in *Car+/+* and *Car-/-* males (supplemental figure 3C). Consistently, publicly available data from a single cell RNAseq study performed on livers from male mice fed a WD for 52 weeks (leading to advanced MASH)^36^ revealed that, both in CD-fed and WD-fed mice, *Car* mRNA expression is almost exclusively located in the hepatocytes (supplemental figure 3A&B).

Altogether, these results demonstrate that CAR is required for the early adaptive transcriptional program of the hepatocytes to the WD, with coordinated shifts in hepatocyte-identity transcription factor target programs consistent with upstream regulation.

### CAR protects from MASH development in a sex-dependent way

We wondered whether this massive transcriptional deregulation in *Car-/-* males could lead to MASH development and fed *Car+/+* and *Car-/-* male and female mice a CD or a WD for 14 weeks (figure 2A). Body weight gain was significantly higher in WD-fed mice (figure 2C), as well as glucose intolerance (supplemental figure 4) and hepatic steatosis (figure 2D-G), but this phenotype was not dependent on the genotype. However, WD-induced liver weight gain and liver damages (illustrated by plasma concentrations of alanine aminotransferase (ALT) and alkaline phosphatase (ALP)) were potentiated by CAR deletion in males (figure 2H&I). WD-induced perturbations of cholesterol metabolism were also significantly higher in *Car-/-* compared to *Car+/+* males, with significantly higher levels of hepatic esterified- and free-cholesterol and of plasma total cholesterol, LDL-cholesterol, total-, primary- and secondary-bile acids, suggestive of bile acid leakage from the liver to the systemic circulation (supplemental figure 5). In females, no significant differences were found between genotypes (figure 2F&G, supplemental figure 5) except slight differences in individual bile acid levels (supplemental figure 5). Interestingly, *Car+/+* and *Car-/-* male and female mice already displayed significantly differences in circulating bile acid levels under CD (supplemental figure 6).

**Figure 2:**
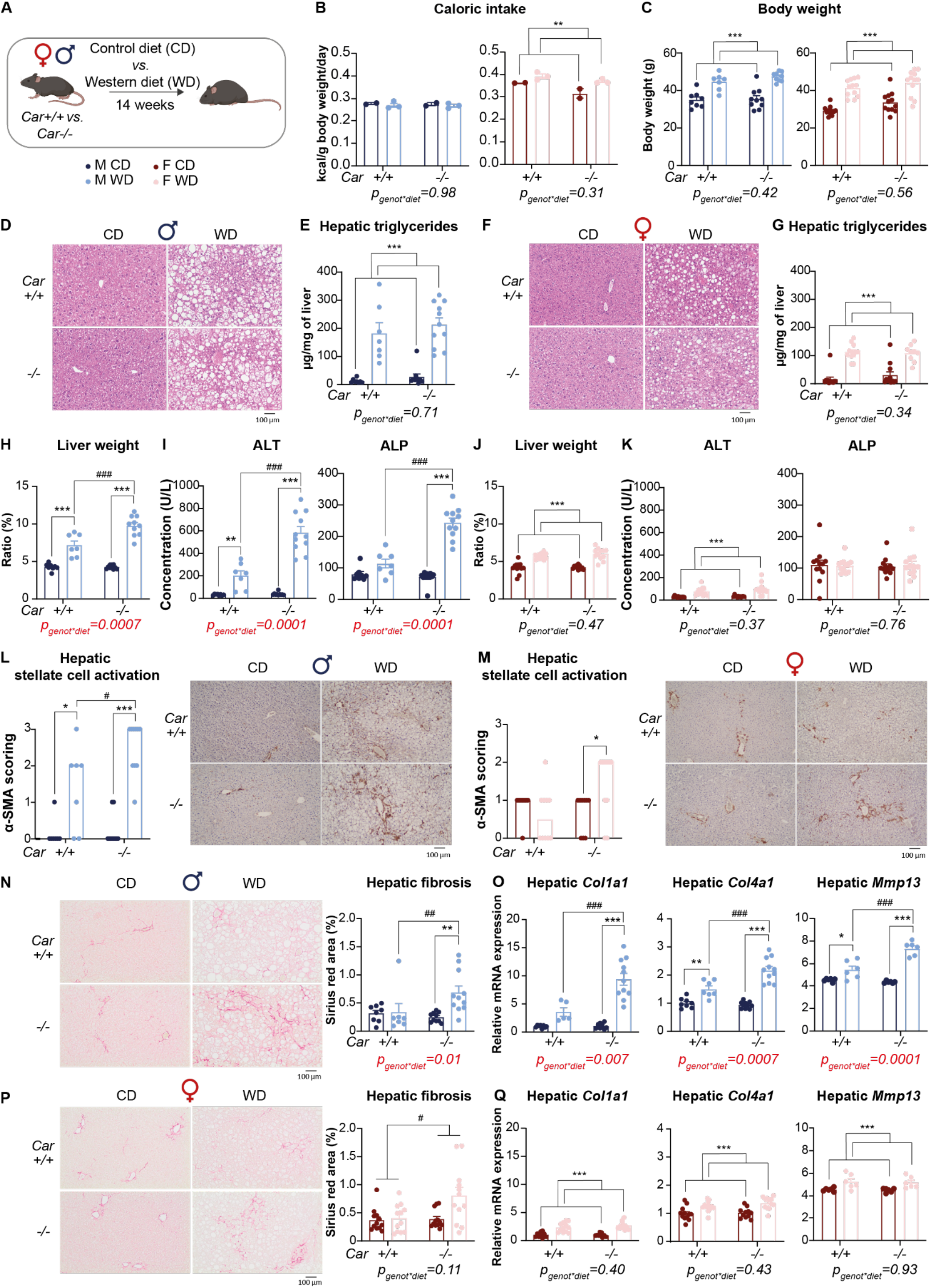
CAR protects male mice from WD-induced MASH. A) Experimental design. B) Mean caloric intake per cage. C) Body weight at the end of the experiment. D) Representative histological sections of male liver stained with haematoxylin and eosin. Magnification x100. E) Male hepatic triglycerides quantified by gas-liquid chromatography. F) Representative histological sections of female liver stained with haematoxylin and eosin. G) Female hepatic triglycerides. H) Ratio of liver to body weight in males. I) Plasma alanine aminotransferase (ALT) and alkaline phosphatase (ALP) concentrations in males. J) Ratio of liver to body weight in females. K) Plasma ALT and ALP concentrations in females. L) Representative histological sections of male liver stained with anti-αSMA antibody. Magnification x100. Stellate cell activation scoring: parenchymal involvement of αSMA staining <3%, 0; 4-33%, 1; 34-66%, 2; >67%, 3. M) Representative histological sections of female liver stained with anti-αSMA antibody and stellate cell activation scoring. Magnification x100. N) Representative histological sections of male liver stained with sirius red. Magnification x100. Quantification of liver fibrosis with sirius red area using ImageJ. O) Hepatic mRNA quantification using RT-qPCR in males. P) Representative histological sections of female liver stained with sirius red. Magnification x100. Quantification of liver fibrosis with sirius red area using ImageJ. Q) Hepatic mRNA quantification using RT-qPCR in females. Results are given as the mean ± SEM. *diet effect, #genotype effect. * or # p<0.05, ** or ## p<0.01, *** or ### p<0.001 (two-way ANOVA). Histological scores were analysed by a non-parametric test (Kruskal-Wallis).

Analysis of the hepatic transcriptome again showed that the number of genes significantly up-regulated in WD-fed compared to CD-fed mice was much higher in *Car-/-* males (1523 DEGs) than in *Car+/+* males (800 DEGs) (supplemental figure 7A). Genes up-regulated in *Car-/-* males specifically were involved in “extracellular matrix organization” and “supramolecular fiber organization”, suggestive of fibrosis development. In females, WD-feeding impacted the hepatic mRNA expression to a much lower extent than in males (supplemental figure 7B). Histological analyses confirmed that WD-induced hepatic stellate cell activation was potentiated by CAR deletion in males (figure 2L), as well as hepatic fibrosis (figure 2N&O), which was observed only in WD-*Car-/-* males. In females, no significant differences were found between *Car-/-* and *Car+/+* (figure 2M, 2P&Q).

We next investigated another nutritional model of hepatic inflammation and fibrosis and fed *Car+/+* and *Car-/-* male mice a Methionine- and Choline-deficient Diet (MCD) (supplemental figure 8). Plasma ALP was higher in MCD-*Car-/-* mice compared to CD-fed mice while no difference was observed between MCD- and CD-*Car+/+* mice. The same was seen for stellate cell activation, consistent with the hepatic mRNA expression of *Ccl2, Adgre1, Col1a1* and *Col3a1*, genes involved in chemotaxis, inflammation-mediated cell adhesion and fibrogenesis. Altogether, CAR deletion aggravated diet-induced MASH in male mice in 2 different nutritional models.

We, and others, described previously that female mice are protected against WD-induced metabolic perturbations^13^, which might prevent us from unravellling a potential role of CAR in female MASH pathogenesis. To circumvent this difficulty, *Car+/+* and *Car-/-* female mice were fed a CD or a WD for 35 weeks (figure 3A). This protocol induced similar levels of hepatic damages in *Car+/+* females than those observed in *Car+/+* males after 14 weeks of WD (figure 3E&2I). At the end of the 35 weeks, body and liver weights, hepatic levels of triglycerides, esterified cholesterol and free cholesterol, hepatic damages, hepatic stellate cell activation and hepatic fibrosis were not different between genotypes (figure 3B-H). This independent long-term study thus confirmed that CAR was not involved in MASH pathogenesis in females.

**Figure 3:**
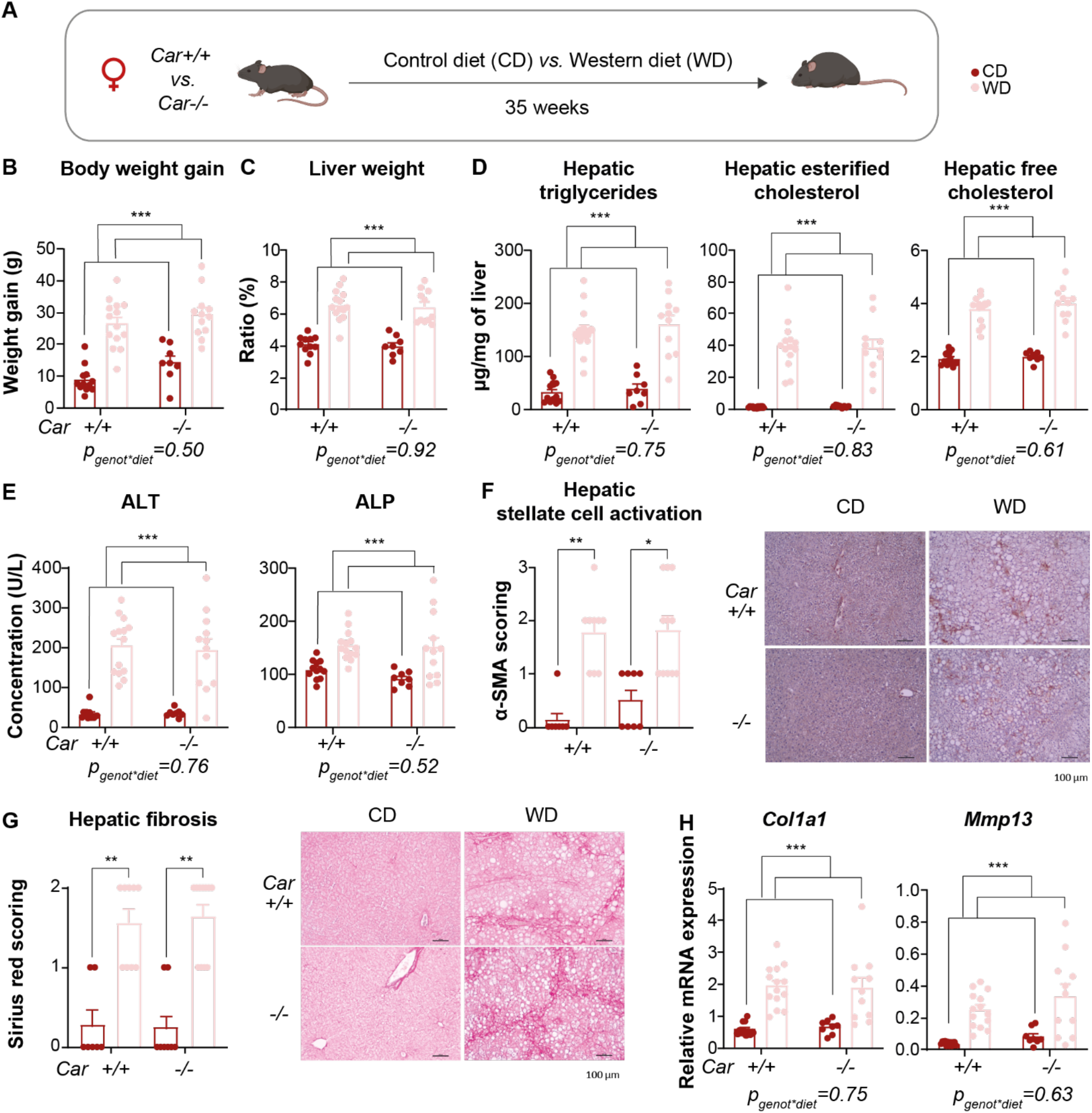
CAR is not involved in WD-induced MASH in females. A) Experimental design. B) Body weight gain. C) Relative liver weight. D) Hepatic triglycerides, esterified cholesterol and free cholesterol quantified by gas-liquid chromatography. E) Plasma alanine aminotransferase (ALT) and alkaline phosphatase (ALP) concentrations. F) Representative liver histological sections stained with anti-αSMA antibody. Magnification x100. Stellate cell activation scoring: parenchymal involvement of αSMA staining <3%, 0; 4-33%, 1; 34-66%, 2; >67%, 3. G) Representative histological sections of liver stained with sirius red in each group. Magnification x100. Liver fibrosis quantified on histological sections. H) Hepatic gene expression assessed by RT-qPCR. Results are given as the mean ± SEM. *diet effect. * p<0.05, ** p<0.01, ***p<0.001 (two-way ANOVA). Histological scores were analysed by a non-parametric test (Kruskal-Wallis).

### A crosstalk between CAR and the gut microbiota influences the hepatic phenotype in males

We next addressed whether the gut microbiota-liver axis could be involved in this male-specific CAR-dependent MASH susceptibility. We first studied the impact of CAR deletion on gut microbiota composition in the WD-experiment. After 14 weeks of WD, relative caecum weight was decreased in both genotypes compared to CD groups (figure 4A), while microbiota α-diversity was not significantly different between groups (figure 4B). As expected, β-diversity was significantly modulated between CD and WD groups in both genotypes (figure 4C). However, a significant shift in the gut microbiota composition was also observed between *Car-/-* and *Car+/+* mice, both under CD (R^2^=0.09, P_adj_=0.0012 using PERMANOVA) and WD (R^2^=0.17, P_adj_=0.002). Bootstraping analysis of the intergroup distances demonstrated that the differences in microbiota composition between *Car+/+* and *Car-/-* mice were significantly greater in WD-than in CD-fed mice (figure 4D), thus demonstrating that pre-existing differences in gut microbiota composition were amplified upon WD.

**Figure 4:**
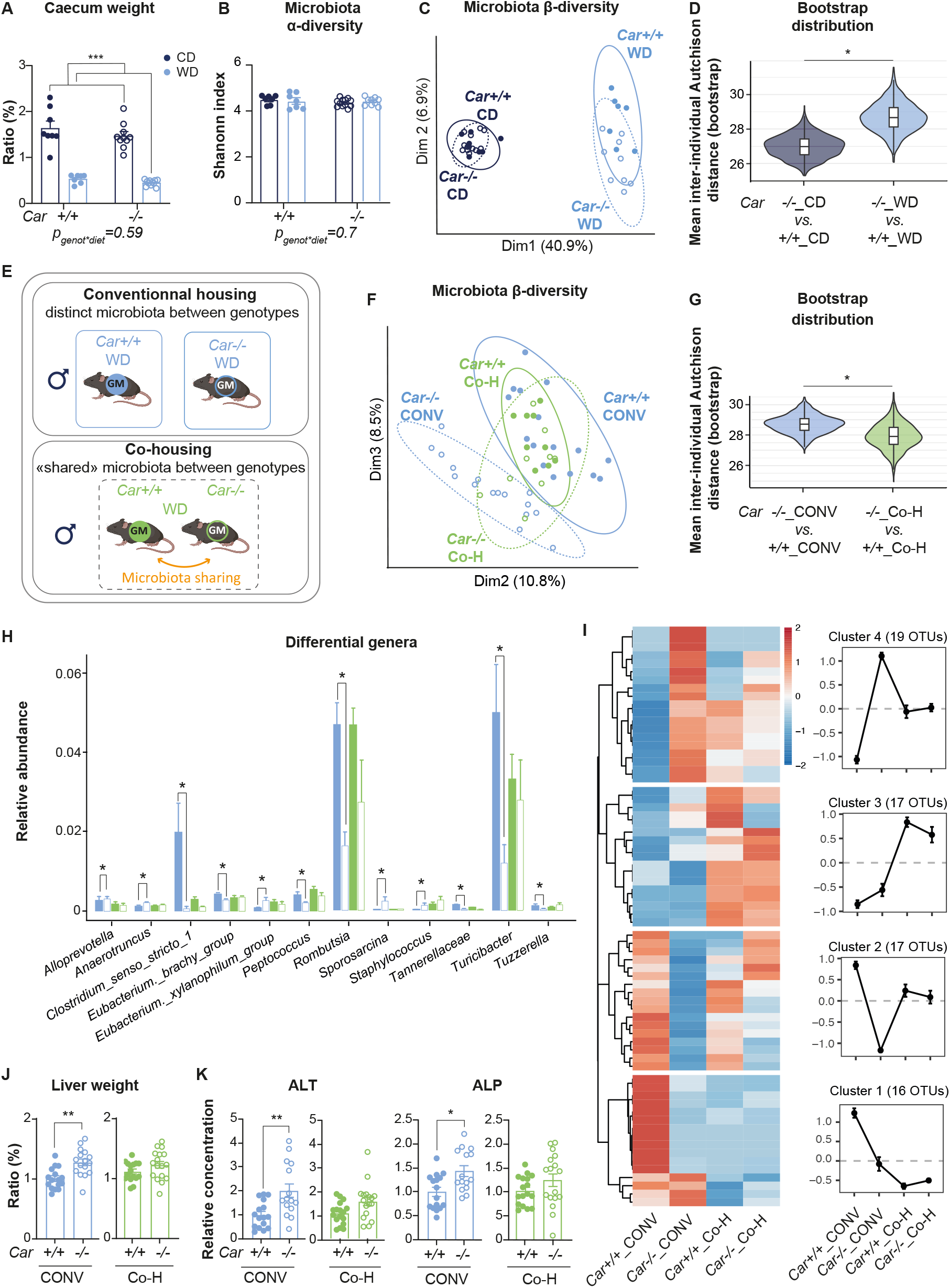
Interactions between CAR and the gut microbiota influence the hepatic phenotype. A) Relative caecum weight in male mice fed CD or WD for 14 weeks. B) Microbiota α-diversity using Shanonn index. C) Principal Component Analysis (PCA) score plots of the microbiota β-diversity using Euclidean distances on CLR-transformed data, equivalent to Aitchison distances. Each dot represents a sample projected onto first (horizontal axis) and second (vertical axis) PCA variables. D) Inter-group distances using the Aitchison distances performed with 1000 bootstrap iterations. E) Experimental design of the co-housing experiment. F) Principal Component Analysis (PCA) score plots of the microbiota β-diversity. G) Inter-group distances. H) Relative abundance of the differentially abundant genera between Car+/+ and Car-/- males identified with Maaslin2. I) Mean relative abundance per group of the differentially abundant OTUs between Car+/+ and Car-/- males identified with Maaslin2. J) Relative liver weights. K) Plasma alanine aminotransferase (ALT) and alkaline phosphatase (ALP) concentrations. Results are given as the mean ± SEM. * p<0.05, ** p<0.01, *** p<0.001 (using two-way ANOVA for 14 weeks WD and unpaired t-tests for the co-housing study).

To further investigate the influence of gut microbiota on CAR hepatic activity, we depleted the gut microbiota in *Car+/+* and *Car-/-* male mice using antibiotics and investigated the CAR-dependent hepatic response using RNA-sequencing (supplemental figure 9A). As previously observed^33^, gut microbiota depletion decreased CAR hepatic transcriptional activity in *Car+/+* mice, as illustrated by lower expression of its main target gene *Cyp2b10* (supplemental figure 9B). Surprisingly, antibiotic treatment modulated a higher number of genes in *Car-/-* males than in *Car+/+* mice (supplemental figure 9C-D). Enrichment analysis revealed that genes upregulated upon antibiotic treatment in *Car-/-* mice specifically were involved in “sterol biosynthetic process” and “lipid localization” (supplemental figure 9E). These results confirmed that CAR integrates microbial signals and responds by modulating the expression of hepatic genes involved in sterol and lipid metabolism.

Next, we hypothesized that this CAR-microbiota dialogue influenced the WD-induced phenotype in *Car-/-* males and performed a co-housing experiment in *Car+/+* and *Car-/-* male mice fed a WD for 14 weeks, allowing gut microbiota sharing by coprophagy between the 2 genotypes (figure 4E). Co-housing led to a similar hybrid gut microbiota composition in both groups, as seen in microbiota β-diversity analyses (figure 4F): PERMANOVA analysis confirmed that the significant difference observed between gut microbiota of conventionally-housed (CONV) *Car+/+ vs. Car-/-* (R^2^=0.13, P_adj_=0.0015) was not seen in co-housed (Co-H)-*Car+/+ vs. Car-/-* (R^2^=0.05, P_adj_=0.35) and bootstrapping analysis confirmed that the inter-group distance was significantly lower between co-H than between CONV mice (figure 4G). At the genus level, relative abundances of *Clostridium sensu stricto 1, Eubacterium brachy group, Peptococcus, Romboutsia, Tannerellaceae, Turicibacter* and *Tuzzerella* were significantly lower and those of *Alloprevotella, Anaerotruncus, Eubacterieum xylanophilum group, Sporosarnica* and *Staphylococcus* were higher in WD-*Car-/-* compared to WD-*Car+/+* males when mice were conventionally housed, but these differences disappeared upon co-housing (figure 4H). Mean relative abundance at the Operational Taxonomic Unit (OTU) level showed 2 clusters (2 and 4 containing 17 and 19 OTUs respectively) with OTUs significantly less (cluster 2) or more (cluster 4) abundant in CONV-*Car-/-* compared to CONV-*Car+/+* that displayed intermediate abundance levels in Co-H mice, thus illustrating that Co-H mice indeed developed a hybrid microbiota (figure 4I). Phenotypically, no significant differences were found between Co-H-*Car+/+* and Co-H-*Car-/-* in WD-induced liver phenotype (figure 4J&K). Thus, CAR-dependent gut microbiota influenced the metabolic phenotype of WD-fed *Car-/-* males.

### Both CAR-dependent microbiota and genotype protect males from WD-induced hepatic damages

We next tested two hypotheses consistent with the observed outcomes: 1) *Car+/+* microbiota could have a beneficial impact into *Car-/-* host or 2) *Car-/-* microbiota undergoes adaptations when residing in a *Car+/+* host, leading to the establishment of a hybrid microbiota beneficial to the metabolic phenotype. To investigate these hypotheses, we performed 2 Fecal Microbiota Transplantation (FMT) studies.

In the first one, *Car-/-* recipient males were transplanted with fecal microbiota from *Car+/+* males (*Car-/-* with Gut Microbiota *Car+/+* (noted *Car-/-* GM*Car+/+)*) and fed a WD for 14 weeks (figure 5A). Fecal microbiota α-diversity was not different between groups (supplemental figure 10A), while β-diversity analysis revealed that gut microbiota composition from *Car-/-* GM*Car+/+* recipient mice was close to that of the *Car+/+* donors (figure 5B&C). Phenotypically, *Car-/-* mice receiving *Car+/+* fecal microbiota (*Car-/-* GM*Car+/+*) displayed lower liver weight and lower hepatic damages compared to *Car-/-* donors (figure 5D&E), thus confirming that the gut microbiota from *Car+/+* mice was protective against WD-induced hepatic perturbations in *Car-/-* mice.

**Figure 5:**
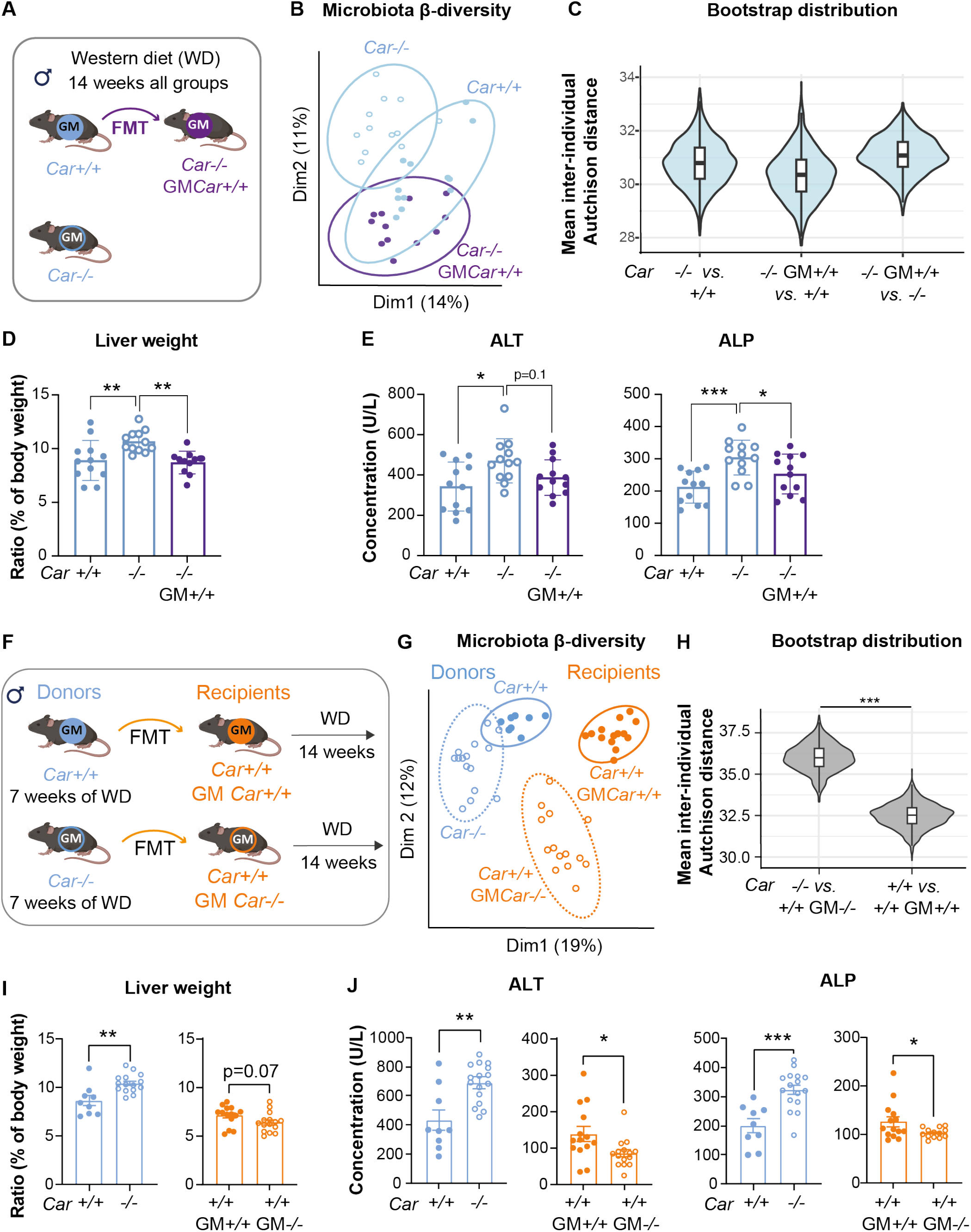
Car+/+ microbiota and Car+/+ genotype protect from WD-induced hepatic damages. A) Experimental design for fecal microbiota transplantation (FMT) study in Car+/+ recipients: Fecal samples from Car+/+ males were transferred into Car-/- recipient mice (Car-/- with Gut Microbiota Car+/+ noted Car-/- GM Car+/+), which were then fed a WD for 14 weeks. B) Principal Component Analysis (PCA) score plots of the microbiota β-diversity using Euclidean distances on CLR-transformed data, equivalent to Aitchison distances. C) Inter-group distances using the Aitchison distances performed with 1000 bootstrap iterations. D) Relative liver weight. E) Plasma alanine aminotransferase and alkaline phosphatase concentrations. F) Experimental design for FMT2 study in Car+/+ recipients: Car+/+ and Car-/- donor males were fed a WD for 7 weeks after which their feces were transferred into Car+/+ recipient mice. All groups were then fed a WD for an additional 14 weeks. G) Principal Component Analysis (PCA) score plots of the microbiota β-diversity using Euclidean distances on CLR-transformed data, equivalent to Aitchison distances. H) Inter-group distances using the Aitchison distances performed with 1000 bootstrap iterations. I) Relative liver weight. J) Plasma alanine aminotransferase and alkaline phosphatase concentrations. Results are given as the mean ± SEM. * p<0.05, ** p<0.01, *** p<0.001 using one-way ANOVA or unpaired Student t-tests.

In the second one, we first aimed to reveal the impact of *Car+/+* host genotype on gut microbiota composition before transplantation and thus fed *Car+/+* and *Car-/-* donors a WD for 7 weeks, then conducted the FMT into *Car+/+* recipients that were subsequently given a WD for 14 weeks (figure 5F). Microbiota α-diversity was not different between groups (supplemental figure 10B), and β-diversity analysis revealed that gut microbiota composition was different between *Car+/+* and *Car-/-* mice in both the donors (PERMANOVA between *Car+/+* and *Car-/-*: R^2^=0.17, P_adj_=0.001) and recipient (R^2^=0.25, P_adj_=0.001) colonies (figure 5G). However, the distance between the *Car-/-* donors and the *Car+/+* recipient from a *Car-/-* microbiota (*Car+/+* GM*Car-/-*) was significantly higher than the distance between the *Car+/+* donors and the *Car+/+* recipient from a *Car+/+* microbiota (*Car+/+* GM*Car+/+*) (figure 5H), illustrating that *Car-/-* microbiota diverged when transplanted into a recipient host expressing CAR. Indeed, a heatmap of the OTU’s significanty differentally abundant between groups showed that the cluster with the highest number of OTUs (54 OTUs in cluster 3, supplemental figure 11) contained OTUs specifically enriched in *Car+/+* GM*Car-/-* but not in the other groups, confirming a significant influence of the recipient genotype in re-shaping the *Car-/-* gut microbiota. At the phenotypic level, *Car+/+* mice receiving *Car-/-* microbiota (*Car+/+* GM*Car-/-*) had significantly lower liver weight and hepatic damages compared to *Car+/+* mice that received a *Car+/+* fecal microbiota (*Car+/+* GM*Car+/+*) (figure 5I-J). Thus, transferring a *Car-/-* gut microbiota to a *Car+/+* host induced compositional shifts that were protective against WD-induced hepatic perturbations, demonstrating that *Car+/+* genotype reshaped the gut microbiota, leading to improved hepatic phenotype.

### Hepatic expression of CAR is decreased in human MASLD

Finally, we tested the human relevance of our findings by investigating the hepatic expression of *CAR* mRNA in different MASLD stages using publicly available gene expression datasets from human cohorts. *CAR* expression was not significantly modulated between healthy individuals and patients with steatosis (figure 6A). However, *CAR* expression was decreased in patients with MASH, cirrhosis or hepatocarcinoma, compared to healthy individuals (figure 6B&C). Next, we compared the networks of the 50 genes showing the highest correlation with *CAR* mRNA expression in human hepatic biopsies diagnosed with mild MASH (F0/2) or advanced MASH (F3/4). Genes significantly correlated with *CAR* expression in patients with mild MASH were no longer significantly associated to *CAR* in patients with severe fibrosis, thus revealing a strong disturbance of *CAR*-associated gene network upon MASH progression (figure 6D). This hepatic expression pattern seemed to be specific to *CAR* since hepatic expression of *PXR* displayed a more erratic pattern (supplemental figure 12), thus showing that the observed decrease in *CAR* expression does not reflect reduced hepatocyte fraction during MASLD progression in these cohorts.

**Figure 6:**
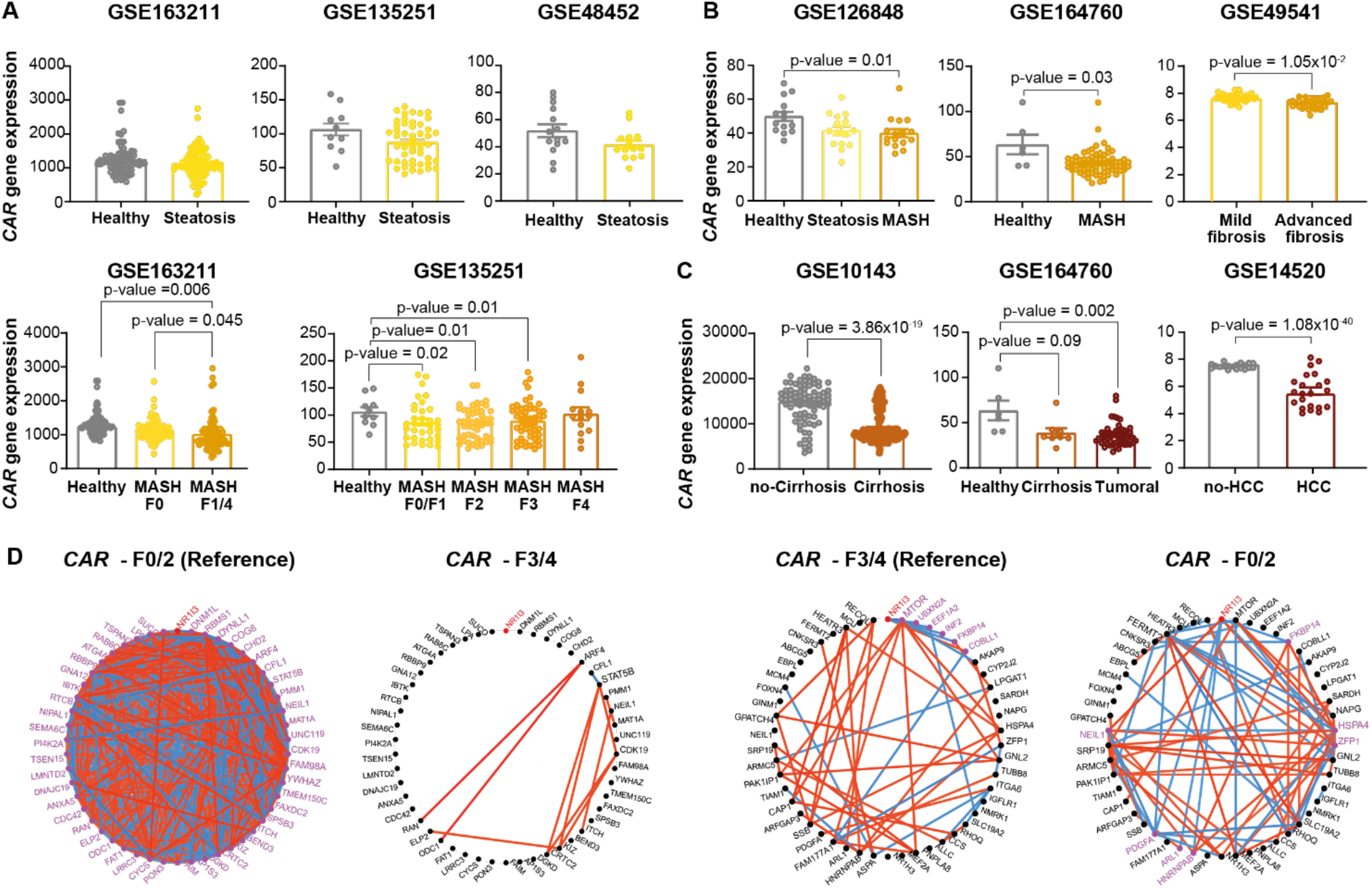
Hepatic expression of CAR mRNA in clinical cohorts of MASLD. (A) CAR mRNA hepatic expression in biopsies from different steatosis clinical cohorts. B) CAR mRNA hepatic expression in biopsies from different MASH clinical cohorts. C) CAR mRNA hepatic expression in biopsies from different cirrhosis/hepatocarcinoma clinical cohorts. D) Networks of the 50 genes showing the highest absolute correlation with CAR mRNA levels in patients with MASH F0/2 and patients with MASH F3/4. The edges corresponding to significant correlations are presented (Bonferroni-adjusted p<0.05). The network circle plot based on the top 50 genes selected in MASH F0/2 is presented on the left while the corresponding circle plot in MASH F3/4 is on the right. Magenta nodes correspond to genes that significantly correlate with the gene of interest. Red and blue were used for positive and negative correlations, respectively.

## DISCUSSION

The gut-liver axis plays a critical role in MASLD pathogenesis, with the local, hepatic, molecular mechanisms connecting dietary, hormonal and microbial cues to hepatic dysfunction remaining undefined. In this study, we identify CAR as a central regulator of hepatic transcription under deleterious diets that also integrates microbiota-derived signals to protect against MASLD in a sexually-dimorphic way.

More specifically, we showed a deregulation of the liver transcriptional response to WD when CAR is absent. First, half of the genes sensitive to WD in *Car+/+* mice were not responsive anymore in *Car-/-* mice, indicating that CAR is required to engage a presumably protective transcriptional response to WD. Second, *Car-/-* mice displayed 2-fold more WD-responsive DEGs than their *Car+/+* counterparts, demonstrating that CAR reshapes which genes respond to WD. This buffering role is best explained by CAR acting as an upstream regulator of a broader transcription factor network. Indeed, several transcription factors central to hepatocyte identity and metabolic adaptation, including HNF4α, STAT3 and STAT5A, are activated by WD in *Car+/+* males but suppressed in *Car-/-* males, suggesting that CAR loss prevents their target genes from being properly modulated. HNF4α in particular orchestrates a large gene expression program, essential to hepatocyte differentiation^37^ and liver function^38^ and has been shown to directly drive the transcription of a broad spectrum of genes related to lipid, glucose, drug, and bile acid metabolism, as well as inflammatory response^39^. In the absence of CAR, disruption of HNF4α is likely to destabilize this program broadly, allowing genes that would normally remain transcriptionally constrained to become responsive to the dietary challenge. Moreover, MASLD is a multicellular pathology where hepatic steatosis begins when lipid input and synthesis exceed disposal in hepatocytes, then persistent lipotoxic stress drives inflammation, cell death, and fibrogenesis in other liver cells^40^. The early kinetics of the CAR-mediated transcriptional deregulation (4 weeks of WD) and our observations that CAR is exclusively expressed in hepatocytes also support a hepatocyte-specific role for CAR in MASLD. However, we acknowledge that we did not directly assess CAR, HNF4α or STAT3/5 chromatin occupancy or activation state and therefore cannot ascribe causality to transcription activity changes. Nevertheless, our results support a model in which CAR shapes the early hepatocyte transcriptional response to WD, with coordinated shifts in hepatocyte-identity transcription factor target programs consistent with upstream regulation.

Phenotypically, in male *Car-/-* mice, this massive transcriptional deregulation upon WD led to the development of hepatic steatosis and later, to MASH with perturbations of cholesterol and bile acid metabolism. Bile acids are potent toxic molecules and their metabolism is finely regulated to avoid their accumulation. Clinical and preclinical studies have shown that disruption of intrahepatic bile acid and cholesterol homeostasis is a major driving force of MASLD progression from steatosis to MASH, cirrhosis, and hepatocarcinoma^41–43^. In humans, we also found that CAR expression was decreased upon MASLD progression in multiple cohorts. Thus, using the WD-induced MASLD model, which is nowadays considered as one of the best models recapitulating MASLD in mice^40,44^, our data support a protective role of CAR against MASLD/MASH development. This clarifies several previous rodent studies that described a controversial impact of CAR deletion in other dietary conditions: CAR deficiency was indeed suggested as beneficial, as it was shown to attenuate fibrosis induced by a MASH-promoting diet^45^ or to normalize the HFD-induced serum triglyceride levels^28^ while others have observed that it could lead to spontaneous obesity, diabetes and steatosis during aging^46^. Our findings also support the multiple positive metabolic benefits previously described upon CAR activation with a pharmacological agonist that mitigated hepatic steatosis, increased glucose tolerance and insulin sensitivity and ameliorated cholesterol homeostasis and hypercholesterolemia^35,47–50^. Recently, a combined CAR agonist/PXR antagonist was even successfully developed and showed improvement in diet-induced hypercholesterolemia^51^. Thus, our data suggest that direct pharmacological testing of CAR activation, alone or in combination with other nuclear receptors, might represent an actionable therapeutic target for MASLD or other metabolic diseases.

However, it is worth noting that none of the phenotypic effects of CAR deletion observed in males were observed in female mice, a result confirmed across multiple independent animal cohorts. Interestingly, most studies that investigated the metabolic role of CAR have used male mice exclusively^28,45,49,50^. Only a couple of studies have compared *Car-/-* to WT females and observed that CAR deletion induced only a slight increase in steatosis upon HFD-feeding^52^ or had no major metabolic impact during aging^46^. Sexual dimorphism is well described upon CAR activation by its pharmacological agonist TCPOBOP: we^35^ and others^53,54^ have shown that CAR activation impacts the liver transcriptome and phenotype more strongly and differently in female than in male mice. The molecular mechanisms of these sex-specific regulations are not well understood but one study has shown that it could relate to basal differences in chromatin states between sexes: DNase hypersensitive sites responsive to the CAR agonist TCPOBOP specifically in male liver were indeed enriched for genomic regions with a basal male bias in chromatin accessibility^53^. Here, we also observed strong differences in the hepatic transcriptomic response of *Car-/-* males *vs.* females to the WD, with much less deregulation observed in *Car-/-* females. Whether this relates to basal chromatin state differences between *Car-/-* male and female mice remains to be studied.

The metabolic mechanisms of this sex dimorphism might also be influenced by the inherent sex-dimorphism of cholesterol metabolism. In humans, bile acid synthesis rate is lower in women and composition of the bile acid pool is different between sexes^55–57^, while in mice, females excrete less fecal bile acid and catabolize less cholesterol through bile acid production than males^58^. CAR activity is also intrinsically linked to sex hormones, with estradiol activating CAR *in vitro*, while androgens inactivate it.^59,60^ CAR also seems to impact circulating sex hormones but evidence is sparse and contradictory, with some authors describing a decrease in androgens (testosterone and androstenedione) in *Car-/-* males^61^, while others found increased testosterone in *Car-/-* males and increased estradiol in *Car-/-* females^46^. MASLD is associated with low androgen levels^62,63^ while estrogens are protective^64^. Thus CAR-mediated impact on sex hormones could also be involved in the male-specific phenotype observed in the present work but this would deserve careful evaluation of the tripartite interplay between hormones-CAR and the gut microbiota. Indeed the gut microbiota can also influence sex hormones^65^, and our present results demonstrate that CAR and the gut microbes are in a complex, bidirectional relationship.

This complex relationship includes i) CAR deletion changing gut microbiota composition, both in a basal- and a WD-induced state. This confirmed previous results that compared *Car-/-* to WT mice and found different fecal gut microbiota compositions^66^; ii) reciprocally, gut microbiota suppression modulating CAR transcriptional activity in the liver, notably decreasing *Cyp2b10* mRNA expression, which we had already observed in germ-free mice^33,67^. Unlike PXR^25^, CAR deletion did not tone down the liver response to microbiota depletion, it amplified it, as illustrated by a much larger number of DEGs sensitive to microbiota depletion in *Car-/-* than in *Car+/+* mice. These results support a more complex model in which, again, CAR acts as a buffer for the liver response to microbial signals. The mediators of the mutual interaction between CAR and the gut microbes probably involve bile acids, which are different between *Car+/+* and *Car-/-* mice^66^ (supplemental figure 6) and can greatly influence gut microbiota composition. Reciprocally, whether gut microbiota-derived bile acids could modulate CAR activity is plausible but controversial as some authors have described several bile acids as transrepressor of CAR activity *in vitro*^68^, while others did not reproduce this *ex vivo* in cultured Teff cells^20^. Other mediators could be indole-containing compounds derived from the microbial metabolism of tryptophan, which have been shown to activate (tryptamine, indole 3-pyruvic acid and indole-3-ethanol) or inhibit (3-methyl indole) CAR *in vitro*^69^, while several microbial-derived diindoles are high affinity CAR ligands, able to modulate CAR-dependent gene expression in human primary hepatocytes and in mouse liver^70^. However, whether these metabolites can reach the liver (probably via the portal vein) at sufficient concentrations to modulate CAR activity *in vivo* is unknown.

We tested the hypothesis that the gut microbiota-CAR interaction could play a role in MASLD in males and gathered many concordant observations supporting this statement: i) in a WD-context, co-housing *Car+/+* and *Car-/-* males led to a hybrid gut microbiota and attenuated the phenotypical differences between genotypes; ii) transferring a *Car+/+* microbiota into a *Car-/-* host decreased WD-induced damages, thus demonstrating that *Car+/+* microbiota was protective against MASLD in *Car-/-* mice. Interestingly, transferring the *Car+/+* microbiota into a *Car-/-* host did not significantly shift the gut microbiota composition, while the opposite FMT (transferring a *Car-/-* microbiota into a *Car+/+* host) significantly changed the gut microbiota composition. Moreover, this second FMT improved the liver phenotype compared to that of *Car+/+* mice receiving *Car+/+* microbiota. These results demonstrated that the expression of CAR itself shifted the gut microbiota toward a more favorable composition. Our findings support a more general concept suggesting that the respective roles of the host factors and the gut microbiota in MASLD pathogenesis cannot be completely distinguished, the host genome affecting gut microbiota toward a more favorable composition to protect against MASLD development. Dissecting the respective role of the host genetics *vs.* microbiota is more advanced in inflammatory bowel disease (IBD) where multiple studies have shown for *e.g.* that the caspase recruitment domain member 9 (Card9) gene contributed to the resolution of intestinal inflammation in mice by regulating gene expression directly in the colon^71^, but *Card9* expression also influenced gut microbiota composition, leading to a favorable production of ligands for the AhR^71,72^ (Aryl hydrocarbon Receptor). CAR is mainly expressed in the liver, and to a lower extent in the small intestine^73^. Recent studies have unveiled its role in the gastro-intestinal tract, where CAR has been shown to protect against bile acid toxicity in the small intestine^20^ and against DSS-induced colonic inflammation^74^. Subsequent studies will thus aim at dissecting the respective role of CAR’s transcriptional activity in hepatocytes or in the intestine and that of CAR-dependent gut microbiota in the protection against MASLD.

A limitation of this study is that all causal evidence for CAR involvement in MASLD stems from loss-of-function genetics and microbiota manipulation. Sufficiency of CAR activation for protection against WD-induced steatosis, inflammation and fibrosis, as well as the translational potential of human-selective CAR agonists remain to be formally demonstrated. Another caveat is that all comparisons were performed in gonad-intact mice carrying a germline deletion of *Car*. Because *Car-/-* mice could display altered levels of circulating androgens and estrogens, the male-restricted phenotype documented here cannot be unambiguously attributed to a cell-intrinsic sex-dimorphic function of hepatic CAR versus a downstream endocrine consequence of lifelong CAR loss. Using adult-onset hepatocyte-specific deletion of *Car* will be required to formally separate these possibilities.

In conclusion, this study identifies CAR as a protective nuclear receptor in male MASLD, both through its host-specific role as transcription factor, and through its ability to regulate the gut microbiota. The exact molecular mechanisms at play probably involve complex relationships between CAR, sex hormones and the gut microbes. Understanding them could help design better drugs against various aspects of metabolic disorders since CAR is gaining popularity as potential drug target.

## Acknowledgements

We thank all members of the EZOP (Experimental Zootechny), UMR Toxalim, Toulouse, France, staff for their careful help with this project. We thank the staff from the Genotoul: Anexplo and We-Met facilities for their help.

