## SupplementalMethodsandFigures for "The constitutive androstane receptor protects against metabolic dysfunction-associated steatotic liver disease through sex specific interactions with the gut microbiota"

**Additional methods and files for:**

### Additional methods

#### Mice and diets

*In vivo* studies were performed in a conventional laboratory animal room following the European Union guidelines for laboratory animal use and care. The current project was approved by an independent ethics committee (CEEA-86 Toxcométhique) under authorization numbers 2019123014045837; 2022041516201224 and 2022090915463602. The animals were treated humanely with due consideration to the alleviation of distress and discomfort. All mice were housed at 22°C ± 2°C on a 12 hours light (ZT0–ZT12) 12 hours dark (ZT12–ZT24) cycle and allowed free access to the diet and tap water. ZT stands for Zeitgeber time; ZT0 is defined as the time when the lights are turned on. *Car-/-* mice (backcrossed on the C57BL/6J background) were engineered in Wei *et al.* laboratory[1] and are bred for 15 years in our animal facility. *Car+/-* mice were bred together and gave birth to true *Car+/+* and *Car-/-* littermate mice, which were then separated by sex and genotype at 4 weeks.

For the 4 weeks Western Diet (WD)-cohort, nine-week-old male and female mice were fed a Western Diet (WD, TD.88137, Inotiv) or a Control Diet (CD, Teklad Global 18% Protein Rodent Diet). Mice were randomly allocated to the different experimental groups: *Car+/+* Female CD (n=8, 2 cages) , *Car-/-* Female CD (n=8, 2 cages), *Car+/+* Female WD (n=8, 2 cages), *Car/-* Female WD (n=8, 2 cages), *Car+/+* Male CD (n=6, 2 cages), *Car-/-* Male CD (n=8, 2 cages), *Car+/+* Male WD (n=7, 2 cages), *Car-/-* Male WD (n=8, 4 cages). Prior to euthanasia, mice were anesthetized with isoflurane and xylazine (2%, 2 mg/kg), then mice were euthanized by cervical dislocation.

For the 14 weeks WD-cohort, nine-week-old male and female mice were fed a Western Diet (WD, TD.88137, Inotiv) or a CD (Teklad Global 18% Protein Rodent Diet). Mice were randomly allocated to the different experimental groups: *Car+/+* Female CD (n=12, 3 cages) , *Car-/-* Female CD (n=12, 3 cages), *Car+/+* Female WD (n=12, 3 cages), *Car/-* Female WD (n=12, 3 cages), *Car+/+* Male CD (n=8, 2 cages), *Car-/-* Male CD (n=10, 3 cages), *Car+/+* Male WD (n=7, 2 cages), *Car-/-* Male WD (n=11, 3 cages). Mice were euthanized by cervical dislocation.

For the 35 weeks (long term) WD-cohort, nine-week-old female mice were fed a WD (TD.88137, Inotiv) or a CD (Teklad Global 18% Protein Rodent Diet) for 35 weeks. Mice were randomly allocated to the different experimental groups: *Car+/+* Female CD (n=12, 3 cages), *Car-/-* Female CD (n=8, 2 cages), *Car+/+* Female WD (n=19, 5 cages), *Car-/-* Female WD (n=16, 4 cages). Prior to euthanasia, mice were anesthetized with isoflurane and xylazine (2%, 2 mg/kg), then mice were euthanized by cervical dislocation.

For the co-housing-cohort, nine-week-old male mice were fed a WD (TD.88137, Inotiv) for 14 weeks. Mice from the conventionally-housed (Conv) groups were separated by genotype at 4-weeks of age, while mice from the co-housed (Co-H) groups were mixed with a 1:1 genotype ratio (2 *Car+/+* mice mixed with 2 *Car-/-* per co-housed cage) at the same age. Mice were randomly allocated to the different experimental groups: *Car+/+* Male Conv (n=16, 4 cages), *Car*$^{-/-}$ Male Conv (n=16, 4 cages), *Car+/+* Male Co-h (n=20, 5 cages), *Car-/-* Male Co-h (n=20, 5 cages). Prior to euthanasia, mice were anesthetized with isoflurane and xylazine (2%, 2 mg/kg), then mice were euthanized by cervical dislocation.

For the first fecal microbiota transplantation (FMT) experiment, one colony of mice was used. The donor mice were composed of nine-week-old *Car+/+* and *Car-/-* male mice fed a (WD) (TD.88137, Inotiv) for 14 weeks. Donor mice were randomly allocated to the different experimental groups: *Car+/+* (n=12, 3 cages), *Car-/-* (n=12, 3 cages). In parallel, eight-week-old *Car-/-* recipient male mice were exposed to antibiotic treatment during 7 days: ampicillin (1 g/L, Euromedex), neomycin (1 g/L, Sigma-Aldrich) and vancomycin (0.5 g/L, MP Biomedicals) dissolved in 0.22µm filtered water and provided *ad libitum* in the animal bottles. The antibiotic solutions were renewed every 2 days. After antibiotic treatment, the recipient colony was fasted for 2 hours and treated with 4 oral gavages of polyethylene glycol (PEG) each 30 minutes apart (1 PEG gavage: 300µL of a solution containing PEG 3350 (77.5 g/L, Sigma-Aldrich) sodium chloride (1.9 g/L, Sigma-Aldrich), sodium sulfate (7.4 g/L, Sigma-Aldrich), potassium chloride (0.98 g/L, Sigma-Aldrich) and sodium bicarbonate (2.2 g/L, Sigma-Aldrich) diluted in 0.22 µm filtered animal facility water). Six hours after the last PEG-gavage, the recipient colony was orally transplanted with fecal microbiota extract from the *Car+/+* WD-fed donor and fed a WD for 14 weeks. During the first three weeks of WD, recipient mice were transplanted with fecal microbiota extract from the *Car+/+* WD-donor colony twice a week, then for the rest of the experiment, mice were transplanted once a week (*Car-/-* GM *Car+/+,* n=12, 3 cages). Prior to euthanasia, mice were anesthetized with isoflurane and xylazine (2%, 2 mg/kg), then mice were euthanized by cervical dislocation.

For the second FMT experiment, two colonies of mice were used. The donor colony was composed of eight-week-old *Car+/+* (n=9, 3 cages) and *Car-/-* (n=15, 4 cages) male mice fed a WD (TD.88137, Inotiv) for 7 weeks before the first fecal microbiota collection and then were fed for 14 additional weeks of WD. In parallel, eight-week-old *Car+/+* recipient male mice were randomly allocated to the different experimental groups: *Car+/+* receiving *Car+/+* FMT (*Car+/+* GM*Car+/+,* n=14, 4 cages), *Car+/+* receiving *Car-/-* FMT (*Car+/+* GM*Car-/-,* n=16, 4 cages) and fecal microbiota transplantation was performed as previously described. Prior to euthanasia, mice were anesthetized with isoflurane and xylazine (2%, 2 mg/kg), then mice were euthanized by cervical dislocation.

For the Methionine- and Choline- Deficient (MCD) diet cohort, twelve-week-old male and female mice were fed a MCD (A02082002B, Research Diets) or a CD (A02082003B, Research Diets) for 10 days. Mice were randomly allocated to the different experimental groups: *Car+/+* Male CD (n=5, 1 cage), *Car-/-* Male CD (n=6, 1 cage), *Car+/+* Male MCD (n=14, 3 cages), *Car-/-* Male MCD (n=14, 3 cages). Prior to euthanasia, mice were anesthetized with isoflurane and xylazine (2%, 2 mg/kg), then mice were euthanized by cervical dislocation.

For the ATB-cohort, nine-week-old male mice were treated with a cocktail of three antibiotics (ampicillin (1g/L, Euromedex), neomycin (1 g/L, Sigma-Aldrich) and vancomycin (0.5 g/L, MP Biomedicals)) dissolved in animal facility water and provided *ad libitum* in the animal bottles for 2 weeks. The antibiotic solutions were renewed every 3 days. Control group (CTRL) had *ad libitum* access to tap water. Mice were randomly allocated to the different experimental groups: *Car+/+* Male CTRL (n=6, 2 cages), *Car-/-* Male CTRL (n=6, 2 cages), *Car+/+* Male ATB (n=7, 2 cages), *Car-/-* Male ATB (n=6, 2 cages). Mice were euthanized by cervical dislocation.

**Oral glucose tolerance test (OGTT)**

OGTT was performed after 12 weeks of diet. Mice were fasted for 6h and received an oral glucose load (2g/kg body weight). Blood glucose was measured at the tail vein using an AccuCheck Performa glucometer (Roche Diagnostics) at -15, 0, 15, 30, 45, 60, 90, and 120 minutes.

**Plasma Analyses**

Alanine aminotransferase, alkaline phosphatase, total cholesterol, high-density lipoprotein (HDL-cholesterol) and low-density lipoprotein (LDL-cholesterol) were determined using an ABX Pentra 400 biochemical analyzer (Anexplo facility, Toulouse, France).

**Plasma bile acid profiling**

Bile acids (BA) were extracted from systemic mouse plasma by precipitation with ice-cold methanol. The separation of BAs was carried out on a Symmetry C18 Luna column (250 mm × 2.1 mm, particle size 5 µm) from Phenomenex (Le Pecq, France). The oven temperature was set at 30 °C. Solvent A was water containing 20 mM ammonium acetate, adjusted to pH 8 and solvent B was acetonitrile. Solvents were delivered at a total flow rate of 500 µl $min^{-1}$. After a 5 min plateau with 28% B, the gradient profile was from 28% B to 90% B linearly in 15 min, followed by a 2 min plateau with 90% B. Column re-equilibration was performed for 4 min. The injection cycle was 26 min. The quantification of 24 different BAs (unconjugated, glyco- and tauro-conjugated BA species within the range of 1.5–2000 $nmol/L^{-1}$) was then performed by high-performance liquid chromatography (UFLC-XR device; Shimadzu, Kyoto, Japan) coupled to tandem mass spectrometry MS/MS (QTRAP 5500 hybrid system, equipped with a Turbo VTM ion source; AB Sciex, Foster City, CA, USA). Instrument control, data acquisition and processing were performed using the associated Analyst 1.5.2 software (AB Sciex, Concord, ON, Canada).

#### Liver neutral lipid analysis

Hepatic lipid contents were determined at the end of experiments as described elsewhere [2]. Briefly, following homogenization of tissue samples in 2:1 (v/v) methanol/EGTA (5 mM), lipids corresponding to an equivalent of 2 mg of tissue were extracted according to Bligh and Dyer in chloroform:methanol:water (2.5:2.5:2.1, v/v/v) in the presence of the internal standards glyceryltrinonadecanoate, stigmasterol, and cholesteryl heptadecanoate (Sigma, Saint-Quentin-Fallavier, France). Triglycerides, free cholesterol, and cholesterol esters were analysed by gas chromatography on a Focus Thermo Electron system using a Zebron-1 Phenomenex fused-silica capillary column (5 m, 0.32 mm i.d., 0.50 µm film thickness).

#### Histological analysis

24 hours paraformaldehyde-fixed, paraffin-embedded liver tissue was sliced into 5-µm sections and stained with hematoxylin and eosin (H&E). Steatosis for mouse liver was evaluated according to [3]. Steatosis was measured depending on the parenchymal involvement by steatosis <5%, grade 0; 6-33%, grade 1; 34-66%, grade 2; >67%, grade 3. Sirius red-stained sections were used for evaluation of fibrosis using imageJ for 14 weeks of western diet. For the WD-long term study sirius red-stained sections (kit Abcam ab150681) were used for evaluation of fibrosis as follows: 0, no fibrosis; 1, pericellular and perivenular fibrosis; and 2, focal bridging fibrosis. Immunohistochemical staining of anti-α-smooth muscle actin (α-SMA) (Abclonal, A17910) was also performed. α-SMA labeling score was established according to Akpolat *et al.* [4] as follows; score 0: absence of labeling or <3% of the periportal region; score 1: between 4 and 33%; score 2: between 34 and 66% and score 3:> 67% of the periportal region. The images were taken by Olympus microscope at 100x magnification.

#### Gene Expression

For hepatic microarray data gene expression profiles were performed at the GeT-TRiX facility (GénoToul, Génopole Toulouse Midi-Pyrénées) using Agilent Sureprint G3 Mouse GE v2 microarrays (8x60K, design 074809) following the manufacturer's instructions. Data acquisition and statistical analysis were performed as previously described.[5] A correction for multiple testing was applied using Benjamini-Hochberg procedure to control the False Discovery Rate (FDR). Probes with |log2FC| > 1 and FDR ≤ 0.05 were considered to be differentially expressed between conditions. The enrichment of Gene Ontology (GO) Biological Processes was evaluated using Metascape.[6] Microarray data are available in the ArrayExpress database (http://www.ebi.ac.uk/arrayexpress) under accession number E-MTAB-17734.

For hepatic RNA-sequencing data gene expression, RNA libraries were constructed from 1000 ng of total RNA at the GeT-TRiX facility (GénoToul, Génopole Toulouse Midi-Pyrénées) using Illumina® Stranded mRNA Prep kit (Illumina, San Diego, CA, USA) following the manufacturer's instructions adapted to produce library sizes compatibles with paired end 150bp read length sequencing. Briefly, mRNAs were captures, then fragmented, converted to double stranded cDNAs and sequencing adaptors were ligated. Libraries were amplified with twelve cycles of PCR. The yield and the quality of the libraries were assessed using D1000 ScreenTapes on TapeStation 4200 (Agilent Technologies, Santa Clara). The libraries were then transferred to GeT-PlaGe facilty (GénoToul, Génopole Toulouse Midi-Pyrénées) for sequencing. The molarity of each library was quantified by qPCR using the Kapa Library Quantification Kit. All libraries were pooled and loaded into one sequencing lane on Illumina NovaSeq 6000 using a 2x150 bp paired-end sequencing mode with NovaSeq 6000 S4 Reagent Kit v1.5. Treatments of sequencing data were performed by the GeT-TRiX facility (GénoToul, Génopole Toulouse Midi-Pyrénées) using the computing resources of genotoul bioinformatics platform Toulouse Midi-Pyrenees (Bioinfo Genotoul). Raw BaseCalls were demultiplexed to generate fastq files using bcl2fastq v2.20.0 software. Raw reads were trimmed of one base for each read ends using cutadapt v2.1, to discard the last low-quality base and the first T base synthetically added in the adapter ligation protocol. Quality of raw and trimmed reads was evaluated using FastQC v0.11.7 software. Clean reads were aligned to Mouse genome reference GRCm39 using STAR v2.7.5a software, including informations of transcripts using associated GRCm39 GTF file. FeatureCounts program from Subread v1.6.0 software was used with strandness information to count the number of reads mapping to exons and summarized to the gene level to generate the count table. Analyses were performed under R v4.0.3, edgeR package and Bioconductor packages. Briefly, raw count table was filtered to eliminate undetected genes using filterByExpr function with min.count=5 and n=3. Then the common, trended and tagwise negative binomial dispersions were estimated with the robust mode of estimateDisp function. A negative binomial generalized log-linear model was fitted to count data with the quasi-likelihood (QL) methods using glmQLFit function to moderate the genewise QL dispersion. Pair-wise comparisons between biological conditions were extracted using specific contrasts with glmQLFtest function to identify differential expression. A correction for multiple testing was applied using Benjamini-Hochberg procedure to control the False Discovery Rate (FDR). Probes with $|\log2FC| > 1$ and FDR $\leq 0.05$ were considered to be differentially expressed between conditions. The enrichment of Gene Ontology (GO) Biological Processes was evaluated using Metascape[6]. RNA-seq data are available in the ArrayExpress database (http://www.ebi.ac.uk/arrayexpress) under accession numbers E-MTAB-17739 and E-MTAB-17740.

For real-time quantitative polymerase chain reaction (RT-qPCR), 2 μg RNA samples were reverse-transcribed using the High-Capacity cDNA Reverse Transcription Kit (Applied Biosystems, Foster City, CA, USA). Additional table 1 presents the SYBR Green assay primers. Amplifications were performed using an ABI Prism 7300 Real-Time PCR System (Applied Biosystems, Foster City, CA, USA). RT-qPCR data were normalized to TATA-box-binding protein (Tbp) mRNA levels.

**GSEA analysis for transcription factor (TF) enrichment**

Gene Set Enrichment Analysis (GSEA) was performed on four diet contrasts (WD vs CD in Car+/+ and Car-/- males and females) using the GSEA function from the clusterProfiler R package [7] (v4.14.0), with 1,000 permutations, gene set size limits of 15–500, and Benjamini-Hochberg FDR correction. Genes were ranked by a z-score combining fold-change and significance: $\text{sign}(\text{logFC}) \times \text{qnorm}(1 - P/2)$. We built a custom gene set database comprising 5,635 sets from three sources: (i) 194 refined liver-specific TF target sets from ChIP-Atlas covering 36 TFs; (ii) 254 TF target sets from MSigDB C3:TFT:GTRD (ChIP-seq-derived, mouse orthologs); and (iii) 44 TF perturbation sets from MSigDB C2:CGP, plus 5,143 pathway sets (Hallmark, Reactome, GO:BP) for FDR calibration. For each TF, a single directionally consistent gene set was used across all contrasts to ensure normalized enrichment scores (NES) signs are comparable. DNA-binding TFs with FDR < 0.05 in at least one contrast were retained (n = 24).

**Cell-type attribution of bulk RNA-seq DEGs.**

To determine whether DEGs originated from hepatocytes or non-parenchymal liver cell types, we anchored gene-level cell-type specificity to a public mouse liver single-cell RNA-seq atlas (GSE166504[8]), comprising 82,168 cells spanning hepatocytes and 12 non-parenchymal populations. We restricted the reference to chow-diet (healthy) cells (25,578 cells across 10 cell types) so that specificity scores reflect normal cell identity rather than disease-shifted expression. For each gene, we computed the Tau (τ) cell-type specificity score [$\tau = \Sigma(1 - \hat{x}_i)/(n-1)$, where $\hat{x}_i$ is the mean expression in cell type i normalized to the maximum across types] and defined cell-type-specific gene sets as genes with $\tau \geq 0.6$ and mean log-normalized expression ≥ 0.1 in their top cell type. DEGs were defined from the bulk liver RNA-seq results at FDR < 0.05 and $|\log_2 FC| > 1$ for the western-diet effect in males. We tested enrichment of each DEG list against per-cell-type-specific gene sets by Fisher's over-representation analysis.

**16S rRNA gene sequencing**

We performed 16S ribosomal RNA (rRNA) gene amplification and sequencing with Illumina MiSeq technology, according to the protocol described by the Earth Microbiome

Project, with slight modifications (www.earthmicrobiome.org/emp-standard-protocols). Briefly, frozen extruded caecal content or feces or fecal suspension samples were mechanically disrupted (bead beating), and DNA was extracted with a PowerSoil-htp kit (QIAGEN). From each DNA sample, the 16S rRNA genes from region V3-V4 were PCR-amplified with a composite forward primer and a reverse primer. The reverse primer contained a unique 12-base barcode, designed with the Golay error-correcting scheme, which was used to tag PCR products from respective samples. The composite forward 515F primer sequence was: 5′-*AATGATACGGCGACCACCGAGATCTACACGCT*XXXXXXXXXXXXTATGGTAATT*GT*GTGY CAGCMGCCGCGGTAA-3′ where the italicized sequence is the 5′ Illumina adaptor, the 12 X sequence is the golay barcode, the bold sequence is the primer pad, the italicized and bold sequence is the primer linker, and the underlined sequence is the conserved bacterial primer 515F. The reverse primer 806R used was 5′-*CAAGCAGAAGACGGCATACGAGAT*AGTCAGCCAG*CC*GGACTACNVGGGTWTCTAAT-3′ where the italicized sequence is the 3′ reverse complement sequence of Illumina adaptor, the bold sequence is the primer pad, the italicized and bold sequence is the primer linker and the underlined sequence is the conserved bacterial primer 806R. PCR reactions consisted of Hot Master PCR mix (Quantabio, Beverly, MA, USA), 0.2 mM of each primer, 10–100 ng template, and reaction conditions were 3 min at 95 °C, followed by 35 cycles of 45 s at 95 °C, 60 s at 50 °C and 90 s at 72 °C on a Biorad thermocycler. PCRs products were quantified using Quant-iT PicoGreen dsDNA assay on a BIOTEK Fluorescence Spectrophotometer and a master DNA pool was generated from the purified products in equimolar ratios. The obtained pool was purified with Ampure magnetic purification beads (Agencourt, Brea, CA, USA), and visualized by gel electrophoresis and then sequenced using an Illumina MiSeq sequencer (paired-end reads, 2 × 250 bp) at the Genom'IC plateform from Cochin Institute.

16S rRNA gene sequences were analyzed with QIIME2 (version 2025.7). Sequences were demultiplexed and quality-filtered with the Dada2 method. We used QIIME2 default parameters to detect and correct Illumina amplicon sequence data, and a table of QIIME2 artifacts was generated. To analyze the taxonomy, we assigned features to operational taxonomic units, according to a 99 % threshold of pairwise identity to the SILVA reference database silva-138-99-nb-classifier. Further statistical analyses were performed in R (version 4.4.2). Raw OTU count tables were first filtered to remove low-abundance and low-prevalence features. OTUs with a total abundance below 0.01% across all samples and those present in fewer than 10% of samples were removed. Filtered OTU counts were transformed using a centered log-ratio (CLR) transformation to account for the compositional nature of microbiome data. Principal component analysis (PCA) was performed on the CLR-transformed data to visualize differences between experimental groups using mixOmics package. Pairwise sample distances were calculated using Euclidean distances on CLR-transformed data, equivalent to Aitchison distances using the vegan package. To assess the significance of differences between groups while accounting for non-independence within cages, pairwise PERMANOVA tests were performed with permutations restricted to within-cage exchanges. Permutation-based p-values were adjusted for multiple testing using the Benjamini-Hochberg procedure. Differentially abundant OTUs between groups were identified using Maaslin2 including cage as random effect. OTUs with q-values < 0.05 were considered significant. A Bootstrap analysis of inter-group distances was applied to quantify the separation between groups with i) individuals resampled with replacement within each group ii) for each bootstrap iteration, the mean inter-individual Euclidean distance (Aitchison distance) between each pair of groups computed iii) this procedure was repeated 1,000 times to generate distributions of inter-group distances iv) empirical bootstrap p-values computed as the proportion of iterations in which the mean distance of one pair exceeded that of another. Unprocessed sequencing data are deposited in the European Nucleotide Archive under accession numbers PRJEB127969, PRJEB128167, PRJEB128170 and PRJEB128171.

**Analysis of publicly available datasets for human cohorts**

Nine independent hepatic gene expression datasets were found on the Gene Expression Omnibus data repository accessed in September 2019. GSE163211[9] (n=318) compared hepatic transcriptome from healthy individuals, patients affected by steatosis, MASH F0 or MASH F1/4. GSE135251[10] (n=216) compared hepatic transcriptome between healthy individuals, patients affected by steatosis or MASH F0 or MASH F1 or MASH F2 or MASH F3 or MASH F4. GSE48452[11] (n=63) analyzed hepatic transcriptome from healthy obese and patients with steatosis. GSE126848[12] (n=57) compared hepatic transcriptome from healthy individuals and patients affected by steatosis or MASH. GSE164760[13] (n=205) compared hepatic transcriptome of healthy individuals and patients with MASH or cirrhosis or hepatocarcinoma from different aetiologies. GSE49541[14] (n=72) contains hepatic transcriptome data from patients with mild-MASH or severe-MASH. GSE10143[15] (n=307) compared the hepatic transcriptome from 307 HCC-patient tumoral and non-tumoral tissue. GSE14520[16] (n=64) compared the hepatic transcriptome from healthy individuals and HCC-patients. GSE159088[17] (n=80) compared hepatic transcriptome from healthy individuals and patients with MASH. Values for *NR1I3* and *NR1I2* gene expression were calculated using the GEO2R tool for microarray data and using GREIN[18] for RNA sequencing data.

Independent publicly available liver single cell RNAseq datasets were exploited with the Single Cell Portal [19]. These datasets came from the study conducted by [20], in which WT mice were fed a WD for 52 weeks (n=4 CD-fed male mice and n=4 WD-fed male).

### Correlation analysis

Correlation matrixes were created with gene expression levels providing from GSE159088[17]. Genes were considered as correlated when expression correlation was found to be >0.8 with a p-value <0.05. These correlated genes were assembled in a correlation network. Transcriptional regulatory network was also built using Trrust (https://www.grnpedia.org/trrust/), a manually curated database of mouse transcriptional regulatory network providing information on the interactions between transcription factors (TFs) and their targets and insights into network hubs, motifs and hierarchical organization. Networks of the 50 genes having the highest absolute correlation (Pearson-correlation) with a transcription factor of interest (red node) were displayed using the R function circlePlot. Only the edges corresponding to significant correlations were represented (Bonferroni-adjusted P value < 0.05). Positive and negative correlations were represented by red and blue edges, respectively.

### Other statistical analyses

All univariate statistical analyses were performed using GraphPad Prism version 10 (GraphPad Software, San Diego, CA). Outliers were identified using the ROUT method. The Kolmogorov–Smirnov test of normality was applied to all data. Two-way ANOVA was performed within each sex (when applicable) using diet (Control Diet or Western) and genotype (*Car+/+* or *Car-/-*) as fitting factors for the models and P-value representing interactions are reported. If *Pgenot*diet* or *Pgenot*treatment* was significant, Sidak's multiple comparisons test was used as post-hoc test to determine which group differed from its appropriate control, otherwise P-values representing the main effects from the ANOVA model (namely diet/treatment or genotype, depending on pre-clinical mice cohort) were reported. P<0.05 was considered as significant. Histological scores were analysed by a non-parametric test (Kruskal-Wallis). In MCD-cohort results have been analysed through one-way ANOVA, because of unbalanced group. In FMT and co-housing cohorts, results have been analysed through unpaired t-tests. Results are given as the mean ± SEM.

### Additional table

| Gene | NCBI Refseq | Forward primer | Reverse primer |
| --- | --- | --- | --- |
| *Adgre1* | NM_001355722 | ACACTTCCCACCCTGGGAC | CTGGACAGGAAGCCTCGTTTA |
| *Car* | NM_009803 | GCTGCAAGGGCTTCTTCAGA | CCTTCCAGCAAACGGACAGA |
| *Ccl2* | NM_011333 | GGTGTCCCAAAGAAGCTGTAGTTTT | AGTTGTAGGTTCTGATCTCATTTGGTT |
| *Col1a1* | NM_007742 | CTCCGGCTCCTGCTCCTC | GGGTTTCCACGTCTCACCAT |
| *Col3a1* | NM_00930 | TGGCACAGCAGTCCAACGTA | CATAGGACTGACCAAGGTGGCT |
| *Col4a1* | NM_009931 | CACGAGGAGCGCAGCC | CAGTCACATTTTCCACAGCCA |
| *Cyp2b10* | NM_009999 | TTTCTGCCCTTCTCAACAGGAA | ATGGACGTGAAGAAAAGGAACAAC |
| *Mmp13* | NM_008607 | ACCTGGACAAGCAGTTCCAAAG | AGCTCATGGGCAGCAACAATAA |

***Additional Table 1:*** *Primers.*

### Additional figures

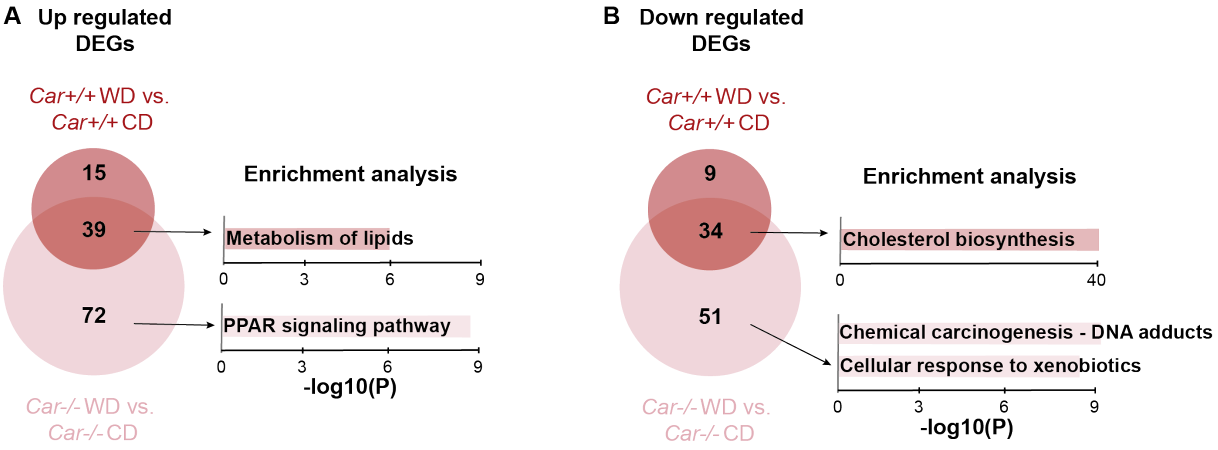


***Supplemental figure 1: Hepatic RNA-sequencing analysis in female mice fed a WD for 4 weeks.*** *A) Venn diagram representing the number of genes significantly upregulated by WD in Car+/+ and Car-/- female mice (FC>2, $p_{adj}$<0.05) and pathway enrichment analysis. B) Venn diagram representing the number of genes significantly downregulated by WD in Car$^{+/+}$ and Car$^{-/-}$ female mice (FC<-2, $p_{adj}$<0.05) and pathway enrichment analysis.*

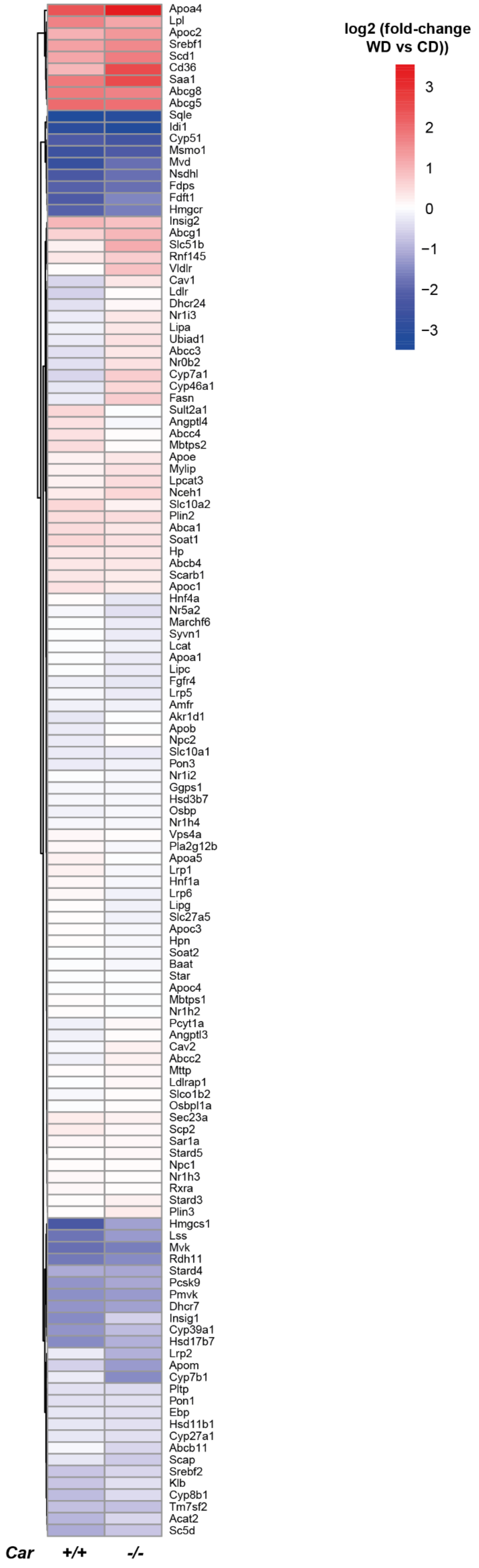


***Supplemental figure 2: Hepatic mRNA expression for genes involved in cholesterol metabolism in male mice fed a WD for 4 weeks.*** *Heatmap representing the log2(fold-change) of gene expression between WD- and CD-fed Car+/+ and Car-/- males for genes involved in cholesterol metabolism.*

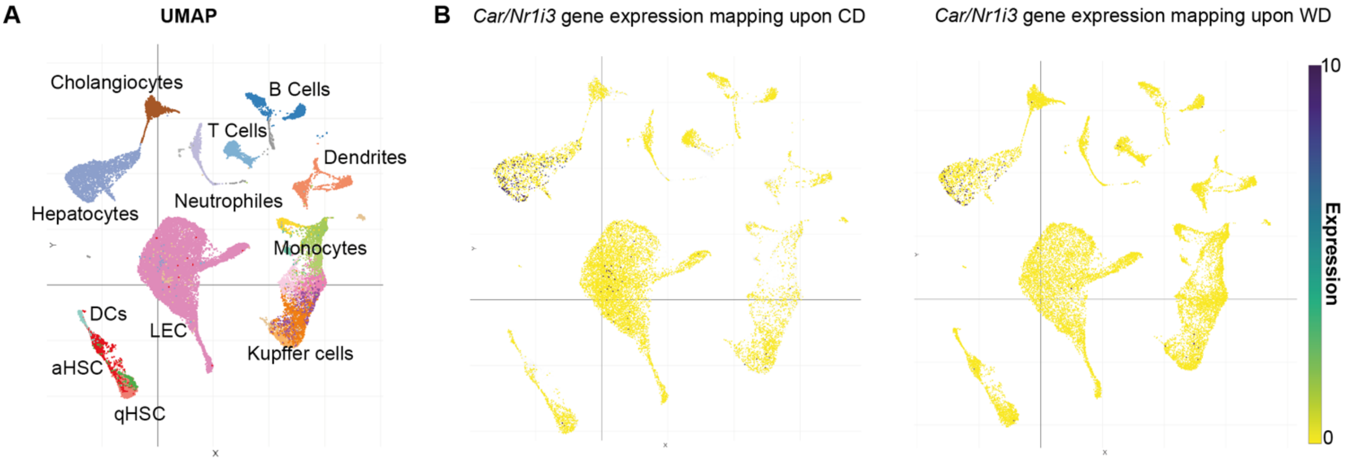


C

| list_name | cell_type | odds_ratio | pvalue | fdr |
|---|---|---|---|---|
| **KOWD_KOCD_down** | **Hepatic_progenitor** | **5,14** | **3,10E-09** | **3,10E-08** |
| **KOWD_KOCD_down** | **Hepatocytes** | **3,92** | **1,37E-06** | **6,83E-06** |
| KOWD_KOCD_down | LSEC | 1,60 | 2,52E-01 | 8,40E-01 |
| KOWD_KOCD_down | B_cells | 0,00 | 1,00E+00 | 1,00E+00 |
| KOWD_KOCD_down | DC | 0,00 | 1,00E+00 | 1,00E+00 |
| KOWD_KOCD_down | Kupffer_cells | 0,90 | 6,55E-01 | 1,00E+00 |
| KOWD_KOCD_down | MoMF | 0,00 | 1,00E+00 | 1,00E+00 |
| KOWD_KOCD_down | Myofibroblasts | 1,01 | 5,42E-01 | 1,00E+00 |
| KOWD_KOCD_down | Stellate_cells | 0,28 | 9,71E-01 | 1,00E+00 |
| KOWD_KOCD_down | T_NK | 0,00 | 1,00E+00 | 1,00E+00 |
| **KOWD_KOCD_up** | **MoMF** | **7,72** | **3,22E-10** | **3,22E-09** |
| **KOWD_KOCD_up** | **DC** | **4,28** | **2,79E-08** | **1,39E-07** |
| KOWD_KOCD_up | Myofibroblasts | 1,34 | 1,07E-01 | 3,56E-01 |
| KOWD_KOCD_up | Hepatic_progenitor | 1,03 | 4,95E-01 | 8,81E-01 |
| KOWD_KOCD_up | Hepatocytes | 0,87 | 7,24E-01 | 8,81E-01 |
| KOWD_KOCD_up | Kupffer_cells | 0,73 | 7,93E-01 | 8,81E-01 |
| KOWD_KOCD_up | LSEC | 0,80 | 7,48E-01 | 8,81E-01 |
| KOWD_KOCD_up | Stellate_cells | 0,82 | 7,47E-01 | 8,81E-01 |
| KOWD_KOCD_up | T_NK | 1,23 | 3,71E-01 | 8,81E-01 |
| KOWD_KOCD_up | B_cells | 0,21 | 9,91E-01 | 9,91E-01 |
| **WTWD_WTCD_down** | **Hepatic_progenitor** | **8,25** | **8,17E-10** | **8,17E-09** |
| **WTWD_WTCD_down** | **Hepatocytes** | **3,34** | **2,13E-03** | **1,07E-02** |
| WTWD_WTCD_down | B_cells | 2,00 | 2,73E-01 | 9,09E-01 |
| WTWD_WTCD_down | LSEC | 1,48 | 4,00E-01 | 1,00E+00 |
| WTWD_WTCD_down | DC | 0,00 | 1,00E+00 | 1,00E+00 |
| WTWD_WTCD_down | Kupffer_cells | 0,83 | 7,02E-01 | 1,00E+00 |
| WTWD_WTCD_down | MoMF | 0,00 | 1,00E+00 | 1,00E+00 |
| WTWD_WTCD_down | Myofibroblasts | 0,45 | 9,34E-01 | 1,00E+00 |
| WTWD_WTCD_down | Stellate_cells | 0,53 | 8,51E-01 | 1,00E+00 |
| WTWD_WTCD_down | T_NK | 0,00 | 1,00E+00 | 1,00E+00 |
| WTWD_WTCD_up | MoMF | 3,45 | 3,37E-02 | 1,68E-01 |
| WTWD_WTCD_up | T_NK | 2,94 | 2,09E-02 | 1,68E-01 |
| WTWD_WTCD_up | DC | 2,23 | 6,25E-02 | 1,76E-01 |
| WTWD_WTCD_up | Hepatocytes | 1,67 | 8,54E-02 | 1,76E-01 |
| WTWD_WTCD_up | Kupffer_cells | 2,20 | 8,79E-02 | 1,76E-01 |
| WTWD_WTCD_up | Hepatic_progenitor | 1,46 | 1,87E-01 | 3,12E-01 |
| WTWD_WTCD_up | Stellate_cells | 1,37 | 3,11E-01 | 4,44E-01 |
| WTWD_WTCD_up | B_cells | 0,00 | 1,00E+00 | 1,00E+00 |
| WTWD_WTCD_up | LSEC | 0,36 | 9,37E-01 | 1,00E+00 |
| WTWD_WTCD_up | Myofibroblasts | 0,69 | 8,57E-01 | 1,00E+00 |

***Supplemental figure 3: CAR expression in mice liver fed a CD or a WD for 52 weeks.*** *A) UMAP clustering of integrated liver single-cell dataset with annotation of cell populations from* [20] *B) Car/Nr1i3 expression mapping derived from single-cell RNA-seq analysis of healthy and advanced MASH livers (n=4 CD-fed male mice, n=4 WD-fed male mice). C) Cell type analysis of the DEGs derived from the bulk RNA-sequencing analysis of liver samples from Car+/+ (WT) or Car-/- (KO) males fed a CD or a WD for 4 weeks (related to Figure 1H&I). Keys: LSEC: liver sinusoidal endothelial cells, DC: denditric cells, MoMF: monocyte-derived macrophages, T_NK: T/natural killer cells.*

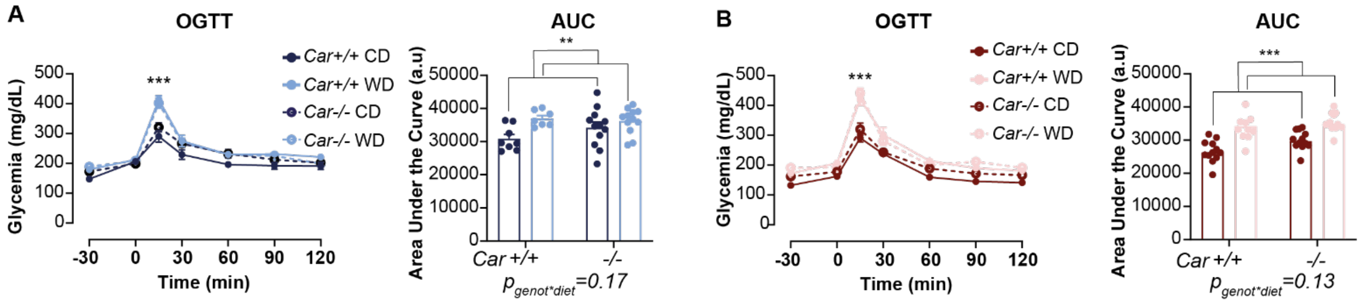


***Supplemental figure 4: Oral glucose tolerance in mice fed a WD for 14 weeks.*** *A) Oral Glucose Tolerance Test (OGTT) assessed after 12 weeks of diet in males. Area Under the Curve (AUC) representing male OGTT results. B) OGTT assessed after 12 weeks of diet in females. AUC representing female OGTT results. Results are given as the mean ± SEM. *diet effect. *p<0.05, ** p<0.01, *** p<0.001 (two-way ANOVA).*

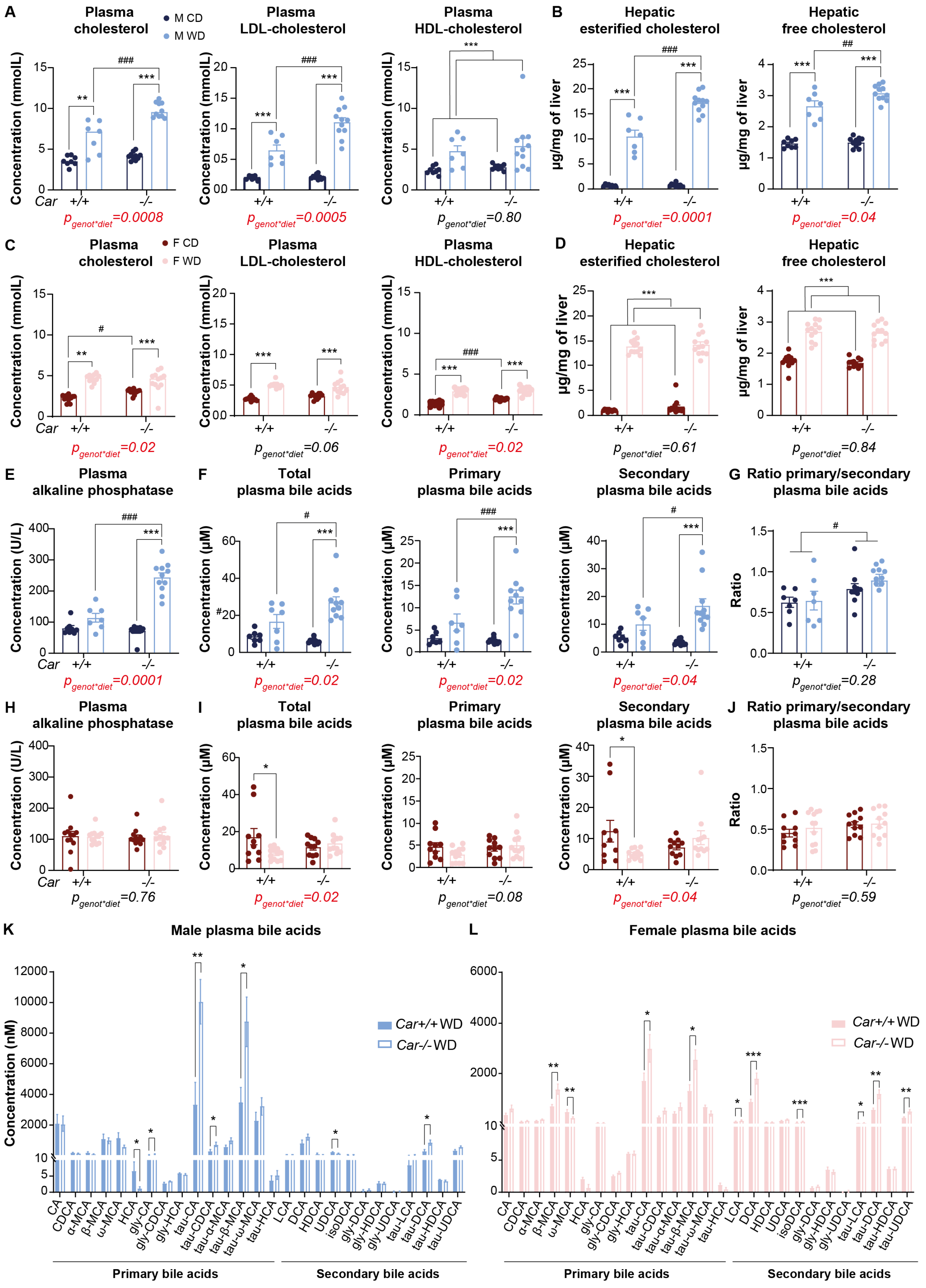
A
Plasma cholesterol
M CD
M WD
Concentration (mmolL)
Car
+/+
-/-
$p_{genot*diet}$=0.0008
Plasma LDL-cholesterol
$p_{genot*diet}$=0.0005
Plasma HDL-cholesterol
$p_{genot*diet}$=0.80
B
Hepatic esterified cholesterol
µg/mg of liver
$p_{genot*diet}$=0.0001
Hepatic free cholesterol
$p_{genot*diet}$=0.04
C
Plasma cholesterol
F CD
F WD
$p_{genot*diet}$=0.02
Plasma LDL-cholesterol
$p_{genot*diet}$=0.06
Plasma HDL-cholesterol
$p_{genot*diet}$=0.02
D
Hepatic esterified cholesterol
$p_{genot*diet}$=0.61
Hepatic free cholesterol
$p_{genot*diet}$=0.84
E
Plasma alkaline phosphatase
Concentration (U/L)
$p_{genot*diet}$=0.0001
F
Total plasma bile acids
Concentration (µM)
$p_{genot*diet}$=0.02
Primary plasma bile acids
$p_{genot*diet}$=0.02
Secondary plasma bile acids
$p_{genot*diet}$=0.04
G
Ratio primary/secondary plasma bile acids
Ratio
$p_{genot*diet}$=0.28
H
Plasma alkaline phosphatase
$p_{genot*diet}$=0.76
I
Total plasma bile acids
$p_{genot*diet}$=0.02
Primary plasma bile acids
$p_{genot*diet}$=0.08
Secondary plasma bile acids
$p_{genot*diet}$=0.04
J
Ratio primary/secondary plasma bile acids
$p_{genot*diet}$=0.59
K
Male plasma bile acids
Concentration (nM)
Car+/+ WD
Car-/- WD
L
Female plasma bile acids
Car+/+ WD
Car-/- WD
CA
CDCA
α-MCA
β-MCA
ω-MCA
HCA
gly-CA
gly-CDCA
gly-HCA
tau-CA
tau-CDCA
tau-α-MCA
tau-β-MCA
tau-ω-MCA
tau-HCA
LCA
DCA
HDCA
UDCA
isoDCA
gly-DCA
gly-HDCA
gly-UDCA
tau-LCA
tau-DCA
tau-HDCA
tau-UDCA
Primary bile acids
Secondary bile acids

***Supplemental figure 5: Cholesterol and bile acid metabolism in mice fed a WD for 14 weeks.*** *A) Male plasma cholesterol, LDL- and HDL-cholesterol concentration. B) Male hepatic esterified and free cholesterol were extracted and analysed by gas-liquid chromatography. C) Female plasma cholesterol, LDL- and HDL-cholesterol concentration. D) Female hepatic esterified and free cholesterol were extracted and analysed by gas-liquid chromatography. E) Male plasma alkaline phosphatase concentration. F) Male plasma bile acids were extracted and analysed by LC-MS. G) Male ratio primary/secondary plasma bile acids concentration. H) Female plasma alkaline phosphatase concentration. I) Female plasma bile acids were extracted and analysed by LC-MS. J) Female ratio primary/secondary plasma bile acids concentration. K) Male plasma bile acids concentration in WD group. L) Female plasma bile acids concentration in WD group. Results are given as the mean ± SEM. *diet effect, #genotype effect. * or # $p<0.05$, ** or ## $p<0.01$, *** or ### $p<0.001$ (two-way ANOVA). Bile acid exhaustive comparisons (panel K and L) were analyzed by multiple t-test comparison.*

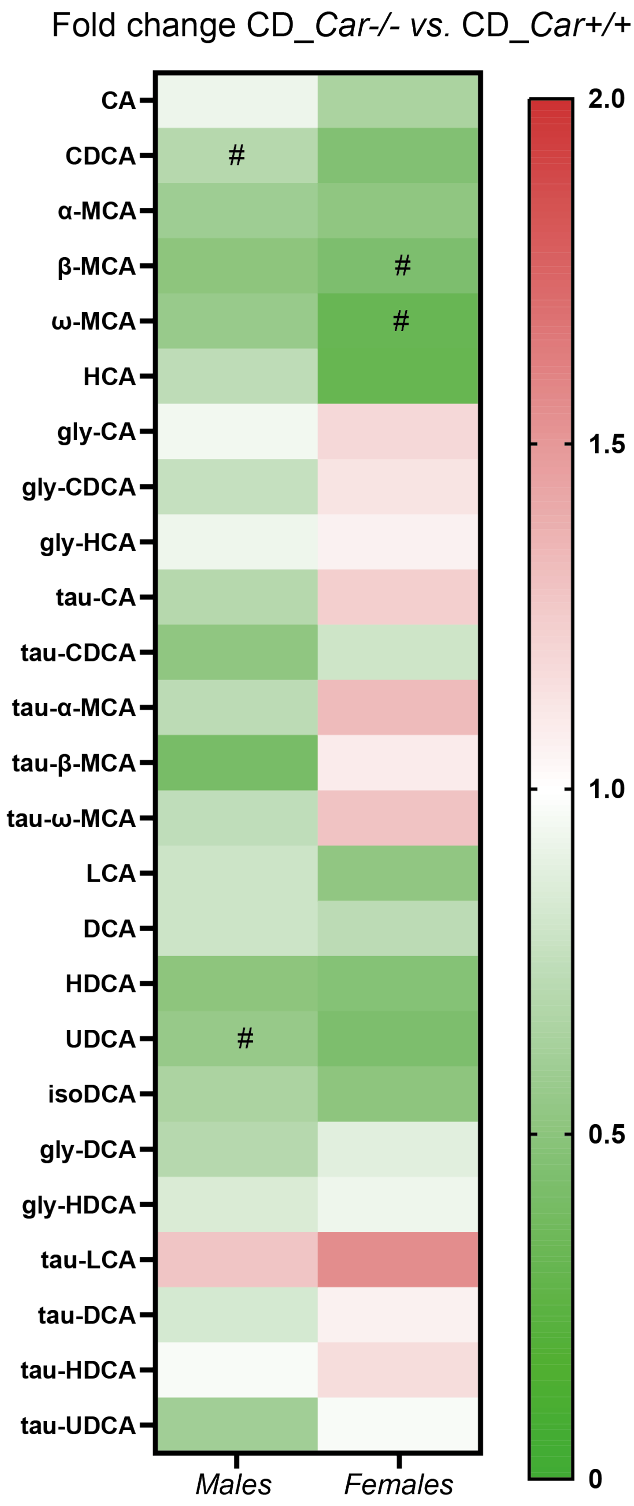


***Supplemental figure 6: Plasma bile acid levels in male and female mice upon a CD.*** *Heatmap representing fold changes of plasma bile acids concentration between Car-/- and Car+/+ in CD group. Results are given as the mean. #genotype effect. # p<0.05 (two-way ANOVA).*

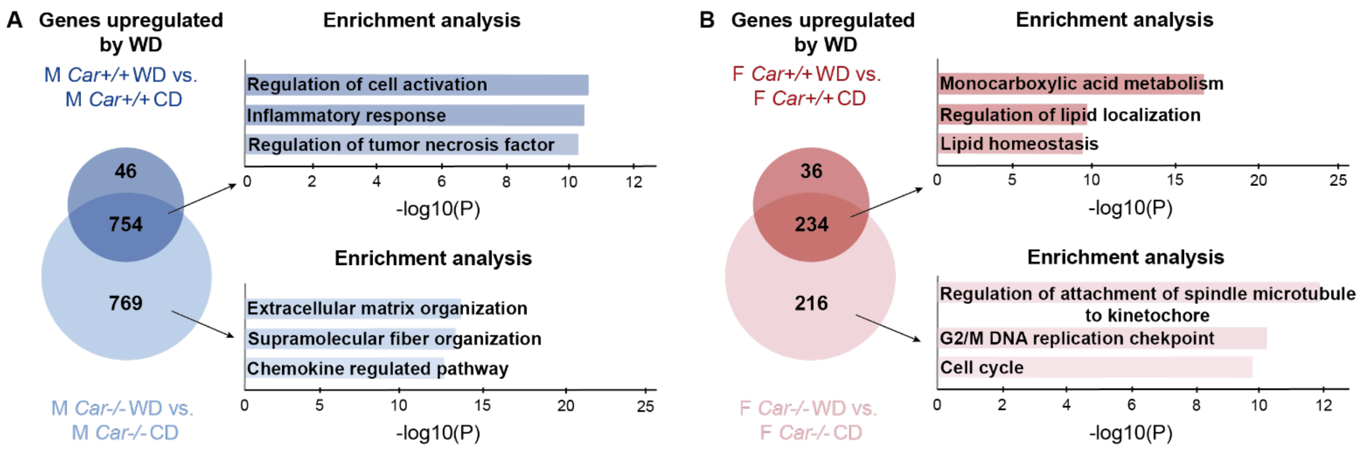


***Supplemental figure 7: DEGs in liver from mice fed a WD for 14 weeks*** *A) Venn diagram representing the number of genes significantly upregulated by WD in Car+/+ and Car-/- female mice (FC>2, padj<0.05) and pathway enrichment analysis. B) Venn diagram representing the number of genes significantly downregulated by WD in Car+/+ and Car-/- female mice (FC<-2, padj<0.05) and pathway enrichment analysis.*

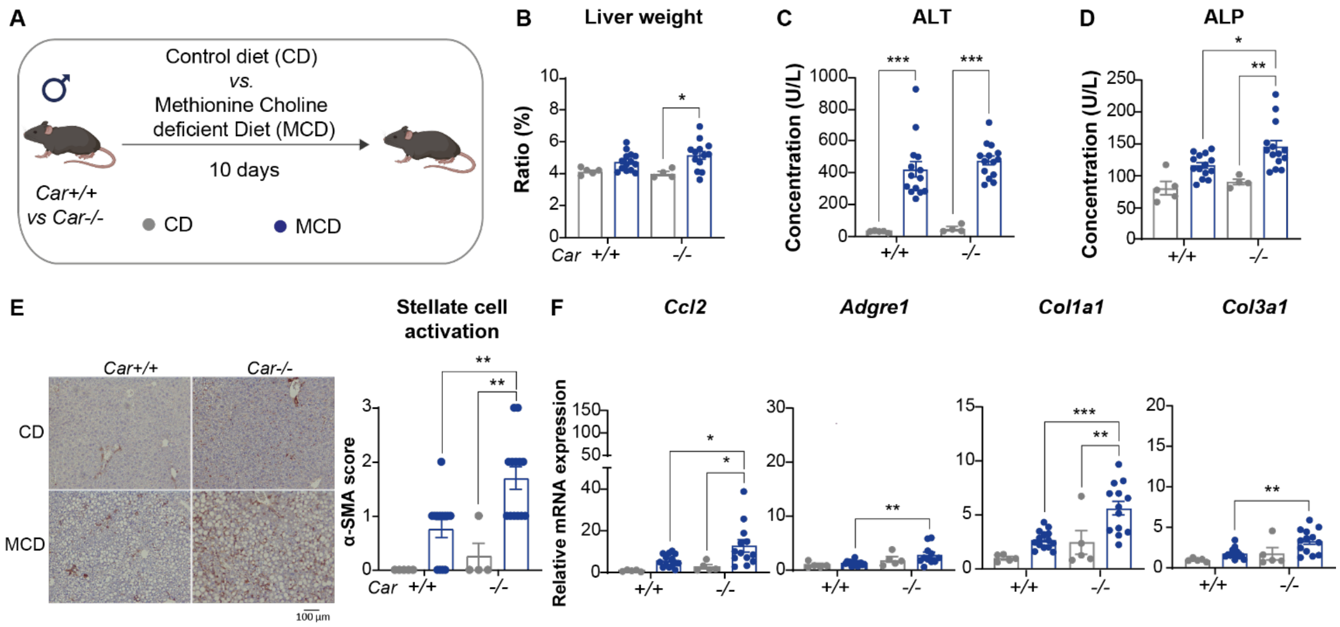


***Supplemental figure 8: Impact of CAR deletion on Methionine Choline deficient Diet (MCD)-induced hepatic fibrosis.*** *A) Experimental design. B) Relative liver weight. C) Plasma alanine aminotransferase concentration. D) Plasma alkaline phosphatase concentration. E) Representative histological sections of liver stained with anti-αSMA antibody in each group. Magnification x100. Male α-SMA sections were used to evaluate hepatic stellate cell activation. Scoring: parenchymal involvement of αSMA staining <3%, 0; 4-33%, 1; 34-66%, 2; >67%, 3. F) Hepatic gene expression assessed by RT-qPCR. Results are given as the mean ± SEM. * p<0.05, ** p<0.01, *** p<0.001 (one-way ANOVA). Histological scores were analysed by a non-parametric test (Kruskal-Wallis).*

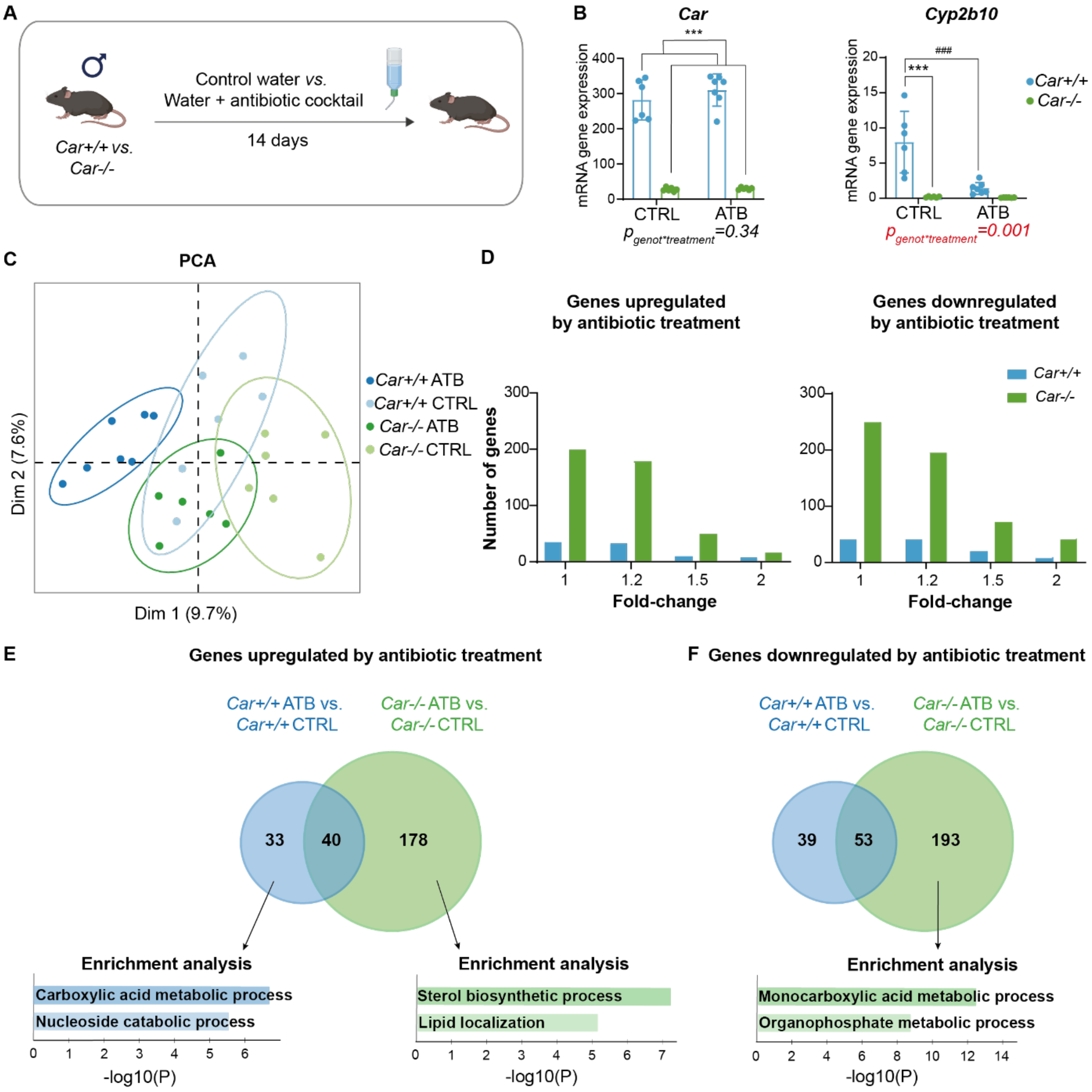


***Supplemental figure 9: Hepatic transcriptome associated to CAR deletion during antibiotic treatment in male mice.*** *A) Experimental design. B) Hepatic gene expression assessed by RT-qPCR. C) Principal component analysis (PCA) score plots of the whole RNA-seq dataset in the liver. Each dot represents an observation (animal) projected onto first (horizontal axis) and second (vertical axis) PCA variables. D) Barplots representing the number of genes significantly upregulated and downregulated by antibiotic (ATB) treatment for each selected fold-change ($p_{adj}$<0.05). E) Venn diagram representing the number of genes significantly upregulated by ATB in Car+/+ and Car-/- mice (FC>1.2, $p_{adj}$<0.05). Enrichment analysis of the 33 genes overexpressed in Car+/+ ATB group. Enrichment analysis of the 178 genes overexpressed in Car-/- ATB group. F) Venn diagram representing the number of genes significantly downregulated by ATB in Car+/+ and Car-/- mice (FC<-1.2, $p_{adj}$<0.05). Enrichment analysis of the 193 genes downregulated in Car-/- ATB group.*

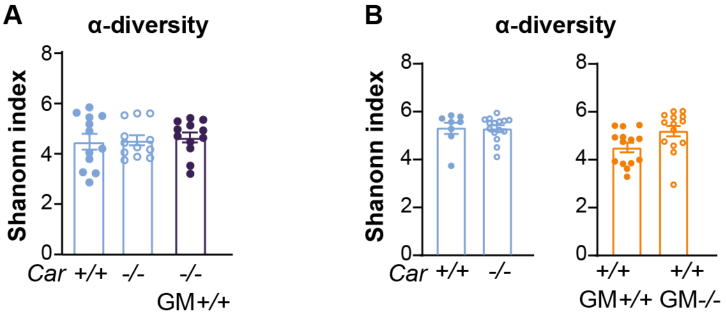


**Supplemental figure 10: Fecal microbiota α-diversity after fecal microbiota transplant experiments.** *A) Fecal microbiota α-diversity using shanonn index in the first experiment of FMT. B) Fecal microbiota α-diversity using shanonn index in the second experiment of FMT.*

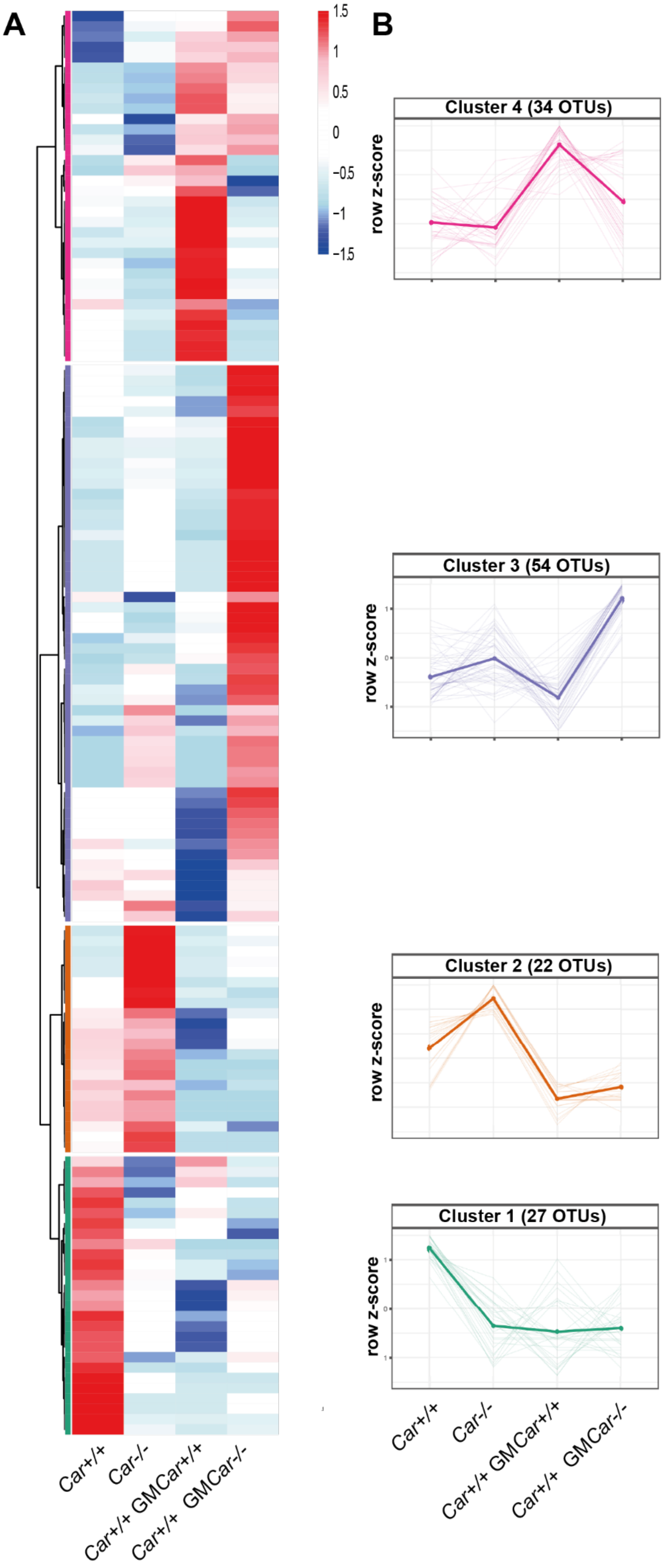

A
B
1.5
1
0.5
0
-0.5
-1
-1.5
Cluster 4 (34 OTUs)
Cluster 3 (54 OTUs)
Cluster 2 (22 OTUs)
Cluster 1 (27 OTUs)
row z-score
Car+/+
Car-/-
Car+/+ GMCar+/+
Car+/+ GMCar-/-

***Supplemental figure 11: Mean OTUs abundance in fecal microbiota of mice from FMT2.*** *A) Heatmap representing the mean OTUs abundance per group identified with Maaslin2. B) Clusters row-z score in each experimental group.*

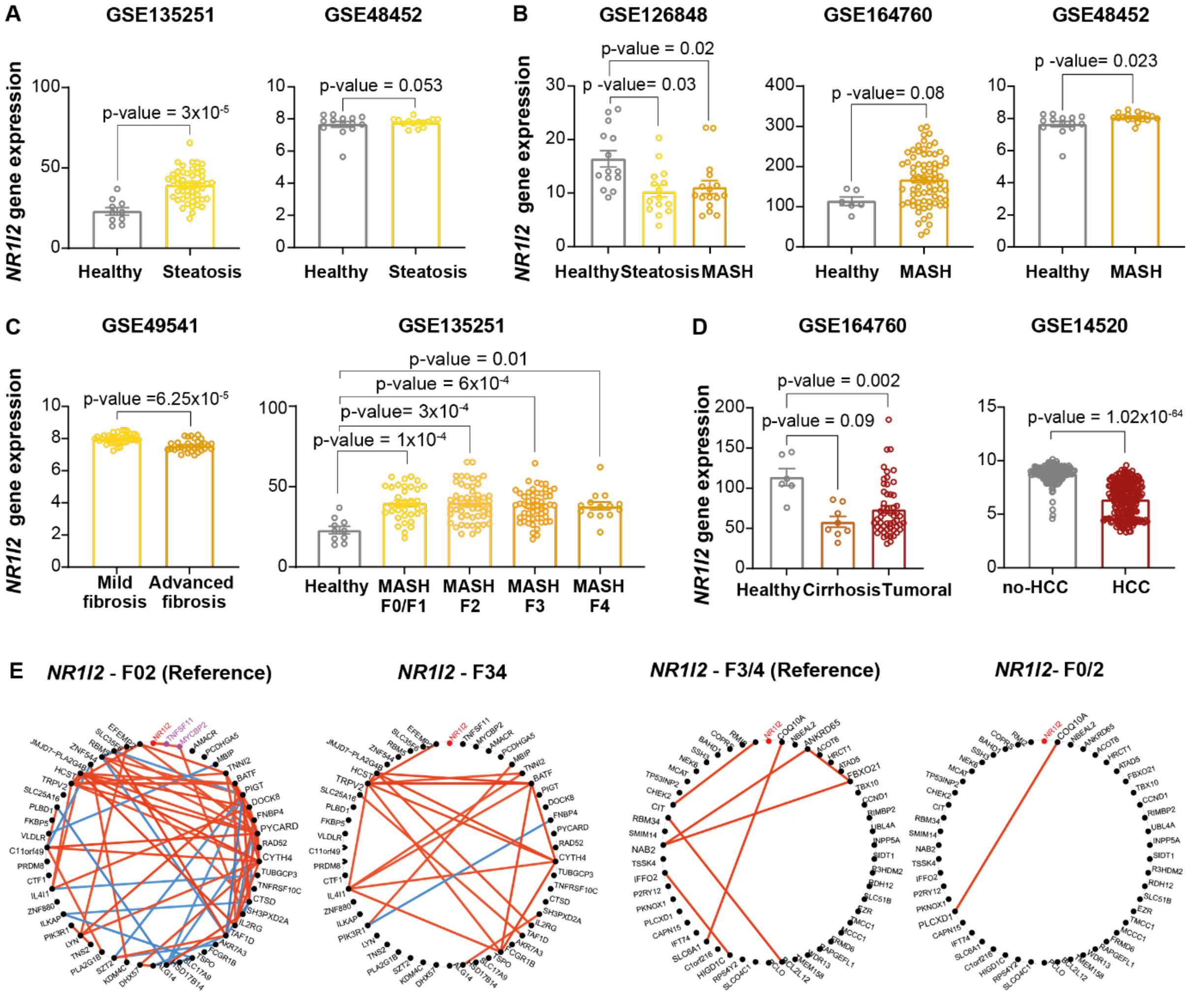


***Supplemental figure 12: PXR (NR1I2) mRNA expression in liver from MASLD human cohorts.*** *A) PXR hepatic expression in biopsies from different steatosis clinical cohorts. B) PXR hepatic expression in biopsies from different MASH clinical cohorts. C) PXR hepatic expression in biopsies from clinical cohorts with different MASH stages. D) PXR hepatic expression in biopsies from different cirrhosis/hepatocarcinoma clinical cohorts. E) Networks of the 50 genes showing the highest absolute correlation with PXR in patients with MASH F0/2 and patients with MASH F3/4. The edges corresponding to significant correlations are presented (Bonferroni-adjusted p<0.05). The network circle plot based on the top 50 genes selected in MASH F0/2 is presented on the left while the corresponding circle plot in MASH F3/4 is on the right. Magenta nodes correspond to genes that significantly correlate with the gene of interest. Red and blue were used for positive and negative correlations, respectively.*
